# Autoencoder and Optimal Transport to Infer Single-Cell Trajectories of Biological Processes

**DOI:** 10.1101/455469

**Authors:** Karren Dai Yang, Karthik Damodaran, Saradha Venkatchalapathy, Ali C. Soylemezoglu, G.V. Shivashankar, Caroline Uhler

**Affiliations:** Department of Electrical Engineering & Computer Science, and Institute for Data, Systems and Society, Massachusetts Institute of Technology, Cambridge, USA; Mechanobiology Institute, National University of Singapore, Singapore; FIRC Institute of Molecular Oncology (IFOM), Milan, Italy

## Abstract

Although we can increasingly image and measure biological processes at single-cell resolution, most assays can only take snapshots from a population of cells in time. Here we describe **ImageAEOT**, which combines an **A**uto**E**ncoder, to map single-cell **Image**s from different cell populations to a common latent space, with the framework of **O**ptimal **T**ransport to infer cellular trajectories. As a proof-of-concept, we apply ImageAEOT to nuclear and chromatin images during the activation of fibroblasts by tumor cells in engineered 3D tissues. We further validate ImageAEOT on chromatin images of various breast cancer cell lines and human tissue samples, thereby linking alterations in chromatin condensation patterns to different stages of tumor progression. Importantly, ImageAEOT can infer the trajectory of a particular cell from one snapshot in time and identify the changing features to provide early biomarkers for developmental and disease progression.

---

Lineage tracing during differentiation, development and disease progression is critical for studying the underlying biological mechanisms. Current experimental methodologies often only provide snapshots of these cellular processes in time and from different cells. Although advances in machine learning in the past decade have prompted new computational methods (1-3), these are primarily aimed at the classification of cells and tissues based on training samples (4-6). This calls for computational methods that infer lineages based on single-cell snapshots in time and identify relevant features of the underlying biological process. Here we present ImageAEOT, which takes advantage of single-cell <u>image</u>s, uses an <u>a</u>uto<u>e</u>ncoder to embed them into a common latent space in order to perform <u>o</u>ptimal <u>t</u>ransport for predicting cellular trajectories from snapshots in time. Such modeling of cellular trajectories allows the identification of functional features and biomarkers underlying a biological process.

The distribution of single-cell chromatin images occupies a low-dimensional manifold in a high-dimensional space (7-8). ImageAEOT models cell trajectories from a source (blue) state to a target (red) state, given examples of images from both populations (Figure 1a). To do this, ImageAEOT first embeds the images into a low-dimensional latent space, which exhibits a simpler geometry than the original data manifold and is learned using a *variational autoencoder* (9-12). Within the latent space, ImageAEOT subsequently learns a model for predicting cell trajectories based on the principle of *optimal transport* (13-16). Finally, ImageAEOT decodes these trajectories back to the image space. Importantly, ImageAEOT does not merely interpolate between the training images, but given an image it infers and generates images of earlier and later cell states. In particular, the generated images resemble the given image, but are endowed with the features of the target population (Figure 1b).

**Figure 1:**
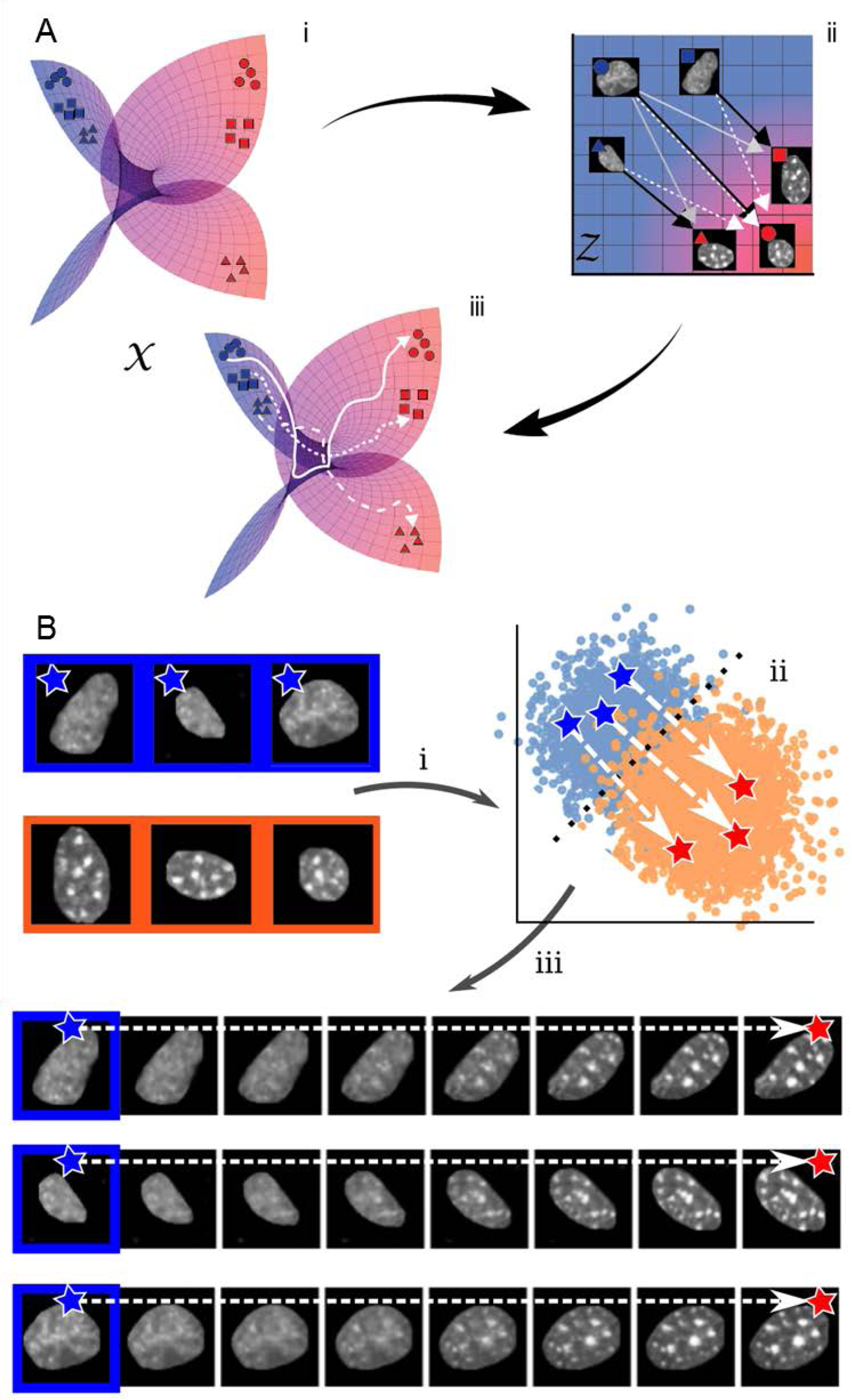
Schematic of ImageAEOT for tracing cell trajectories. (a) *X* represents the manifold of images taken from a population of cells during the biological process of interest. The objective is to trace a trajectory from the blue state, i.e., source population, to the red state, i.e. target population. First, the images are mapped to a latent space *Z* using a variational autoencoder. In the latent space, optimal transport methods are used to trace trajectories from the source population (blue) to the target population (red). Finally, using the variational autoencoder the latent space trajectories are mapped back to the image space, which can be visualized as smooth trajectories from any given image in the source population to a generated image in the target distribution or vice-versa. (b) Illustration of ImageAEOT using nuclear images of MCF7 cells (source population) and NIH3T3 cells (target population). The end points of the predicted trajectories are generated images, i.e. ImageAEOT does not merely interpolate between two given images but rather generate nuclei that have the features of nuclei in the target population, but still resemble the given cell nucleus in the source population.

As a proof-of-concept, we apply ImageAEOT to nuclear and chromatin images during the activation of fibroblasts by tumor cells in engineered 3d tissues (Figure 2a). Cancer-associated fibroblasts play a critical role in tumor initiation and progression (17-19). To understand which fibroblasts get activated within the heterogeneous tissue microenvironment, we apply ImageAEOT to trace the lineages of fibroblasts in a fibroblast-tumor co-culture system (Supp. Figures 1-2). The data consists of single-cell nuclear images of NIH3T3 fibroblasts co-cultured with MCF7 cells for 1-4 days. By Day 4, a subset of fibroblasts are activated and ImageAEOT infers the state of these cells in earlier time points.

**Figure 2:**
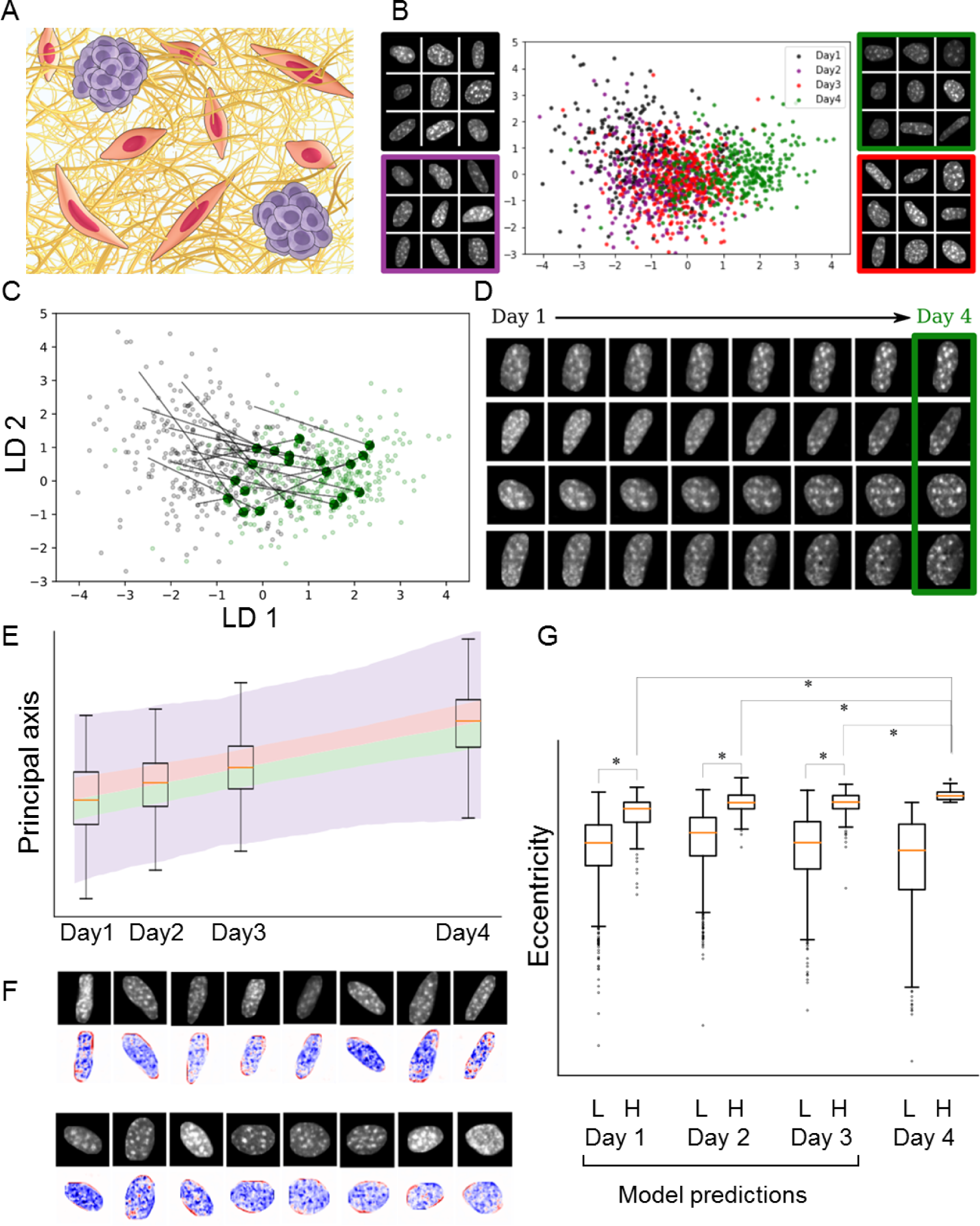
ImageAEOT applied to tracing trajectories of fibroblast activation in a fibroblast-tumor coculture system. (a) Schematic of the fibroblast-tumor coculture experiment. NIH3T3 fibroblasts were cocultured with MCF7 cells for 1 to 4 days. The cells were subsequently fixed, DAPI stained, and imaged. (b) Visualization of NIH3T3 nuclei from Days 1-4 in both the original image space and the latent space using an LDA plot. The first two linear discriminants capture the smooth progression of the cells from Day 1 through Day 4. Day 1: black; Day 2: purple; Day 3: red; Day 4: green. (c) Predicted trajectories in the latent space using optimal transport. ImageAEOT was used to trace the trajectories of Day 4 NIH3T3 nuclei back to Day 1 NIH3T3 nuclei. Each black arrow is an example of such a trajectory. (d) Predicted trajectories mapped back to the image space. Note that only the last image in each sequence is a real Day 4 NIH3T3 nucleus. All other images are predicted and generated by ImageAEOT. (e) Box plot of the principal feature along the first linear discriminant as predicted by ImageAEOT is shown in color (the line separating pink and green is the median; the lines separating green from purple and red from purple denote respectively the first and third quartiles; the blue extends to 1.5 times the interquartile range); the box-plots of the observed experimental distributions are overlaid on top. Note that the distributions of the predicted trajectories coincide with the true distributions, even though only Day 1 and Day 4 NIH3T3 nuclei were used to trace the trajectories. (f) Visualization of the principal feature along the first linear discriminant from (e). The nuclear images are of Day 1 NIH3T3 fibroblasts. The images below show the difference between the generated images along the first linear discriminant and the original image (blue: decrease in pixel intensity; red: increase in pixel intensity). These results suggest that the elongated nuclei become more elongated (i.e. increase of intensity at the poles) and the more spherical nuclei remain more spherical as the fibroblasts progress from Day 1 to Day 4 during their activation. In addition, fibroblast activation is accompanied by chromatin decondensation as revealed by the decrease in pixel intensities. (g) Quantification of subpopulations of NIH3T3 nuclei on Day 4 based on nuclear elongation (L: low eccentricity, H: high eccentricity) back-traced to Days 1-3. The results suggest that a subset of the Day 1 population is already primed for activation and can be identified by ImageAEOT.

Figure 2b shows the low-dimensional latent representation of the NIH3T3 nuclear images obtained by the autoencoder. The linear discriminant analysis (LDA) plot captures the progression of nuclear morphology and chromatin condensation patterns during fibroblast activation. The architecture of our deep convolutional variational autoencoder is shown in Supp. Figure 3. The model was tuned to ensure that images can be encoded and decoded with high fidelity, while also maintaining the geometry of the low-dimensional image manifold in the latent space (Supp. Figures 4-5). As a consequence, classifying cell states using deep convolutional neural networks based on their latent representation achieves a comparable level of accuracy as when classifying cells based on the original images (Supp. Figures 6-9, 14a).

Subsequently, we back-traced NIH3T3 cells from Day 4 to Day 1 in the latent space (Figure 2c) and generated the corresponding single-cell image trajectories using ImageAEOT (Figure 2d). Note that only the last image in each sequence is real; the others are generated. Figure 2e shows that the predicted distributions of Day 2 and Day 3 NIH3T3 nuclei correspond to the observed experimental distributions.

ImageAEOT can also be used to identify biomarkers of the fibroblast activation process by adding small perturbations along the first linear discriminant in the latent space and comparing the corresponding single-cell images in the original space. As shown in Figure 2f, the fibroblast activation process involves an increase in nuclear elongation and alterations in chromatin condensation patterns. Importantly, this analysis reveals that fibroblast subpopulations that are more elongated in Day 1 become more activated in Day 4 (Figure 2g), suggesting that ImageAEOT can identify subpopulations of fibroblasts in the heterogeneous tissue microenvironment that are primed for activation. A similar analysis was performed on the MCF7 nuclei during the activation process (Supp. Figures 10-13, 14b).

Next, we applied ImageAEOT to model trajectories of nuclei progressing through various stages of breast cancer (Figure 3a). The dataset consists of nuclear images of HME-1 (normal breast epithelial) cells, MCF10A (fibrocystic epithelial) cells, MCF7 (metastatic breast cancer) cells, and MDA-MB231 (highly invasive metastatic breast cancer) cells. As described in the earlier sections, the first step of ImageAEOT involves learning a latent representation of the nuclear images that captures their low-dimensional structure (Figure 3b). The quality of the variational autoencoder is evidenced by accurate reconstruction, sampling, and classification results (Supp. Figures 15-19). ImageAEOT was then used to back-trace the trajectories of nuclei from the highly invasive state (MDA-MB231) to normal breast epithelial state (HME-1) in the latent space (Figure 3c). Decoded to the image space, these trajectories yield predictions of how normal mammary epithelial HME-1 cell nuclei may progress through fibrocystic or metastatic stages of breast cancer (Figure 3d). We also analyzed the principal nuclear features that change between the various cell types by adding small perturbations to the latent representations and decoding them to the image space. This analysis shows that the principal features of the transition between MCF10A and MCF7 involve both alterations in nuclear morphology and chromatin condensation patterns (Figures 3e-f), while the transition between other pairs of cell types (HME-1-to-MCF10A and MCF7-to-MDA-MB231) are mainly characterized by nuclear morphological changes (Supp. Figures 20-21).

**Figure 3:**
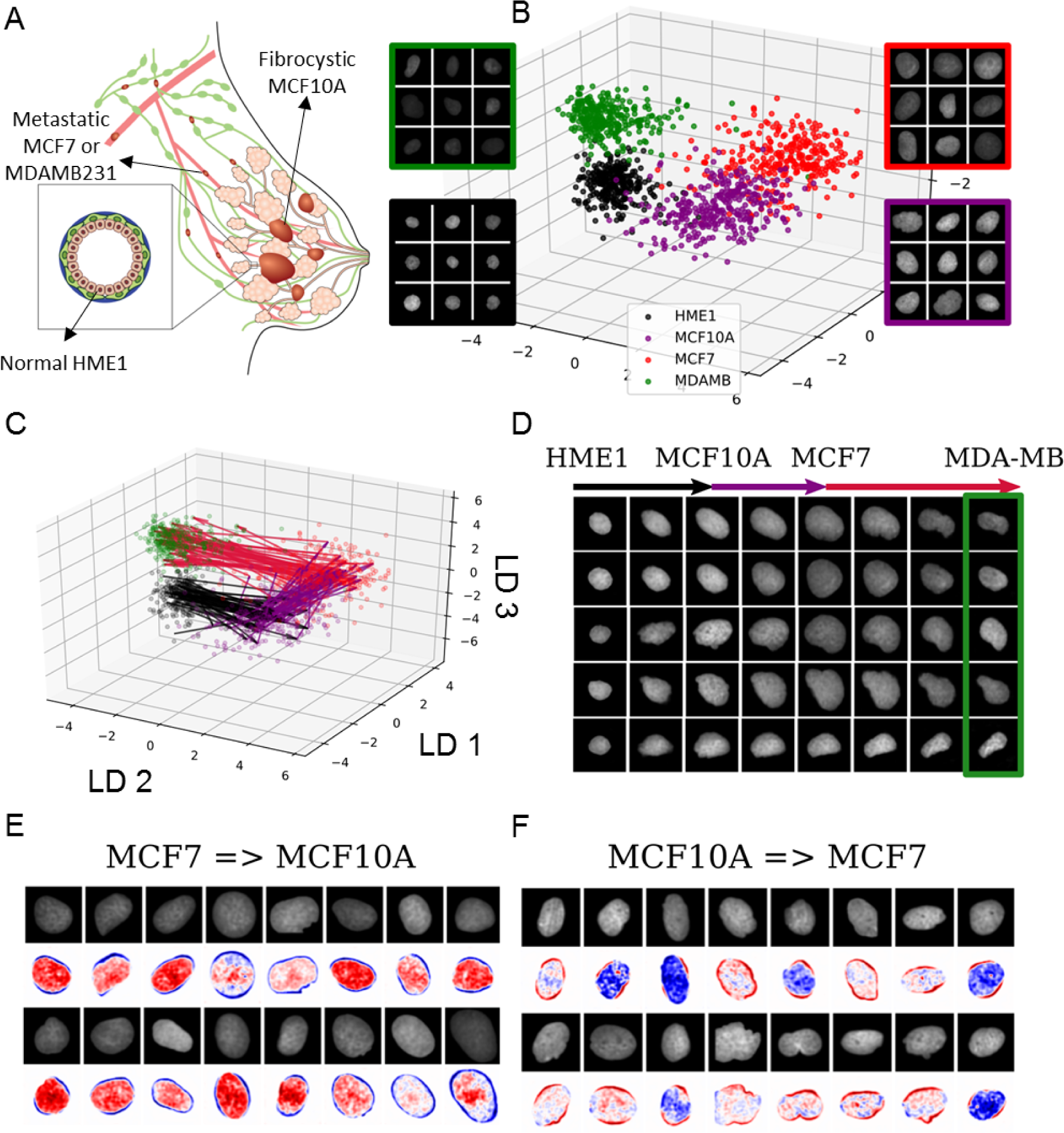
ImageAEOT applied to tracing cellular trajectories during breast cancer progression. (a) Schematic of breast cancer progression. Normal epithelial cells (HME-1) in the breast may become fibrocystic (MCF10A), develop into cancer cells (MCF7), and finally become highly invasive (MDA-MB231). (b) Visualization of nuclear images from four breast cell lines in both the original image space and the latent space using an LDA plot. HME-1: black; MCF10A: purple; MCF7: red; MDA-MB231: green. (c) Predicted trajectories in the latent space using optimal transport. ImageAEOT was used to trace the trajectories from HME-1 to MCF10A to MCF7 to MDA-MB231. (d) Predicted trajectories mapped back to the image space. Note that only the final image in each sequence is a real MDA-MB nucleus; the remaining images are predicted and generated by ImageAEOT. (e-f) Illustration of the principal features that change between MCF10A and MCF7, namely a combination of nuclear morphological and chromatin condensation features.

Finally, we applied ImageAEOT to model trajectories of nuclei in human breast tissues. The dataset consists of nuclear images from normal breast epithelial tissue as well as breast cancer tissue. ImageAEOT learns a high-quality latent representation of these images (Figure 4a), indicated by our reconstruction and sampling results (Supp. Figure 22). The trajectories of nuclei from normal to cancerous cells in the latent space are shown in Figure 4b and the decoded images in Figure 4c. Within the tissue microenvironment the principal features corresponding to the transition between normal and cancer cells involve both alterations in nuclear morphology and chromatin condensation patterns (Figures 4d-e). Collectively, these results suggest that ImageAEOT can be applied to identify early physical biomarkers of cancer initiation and progression (20).

**Figure 4:**
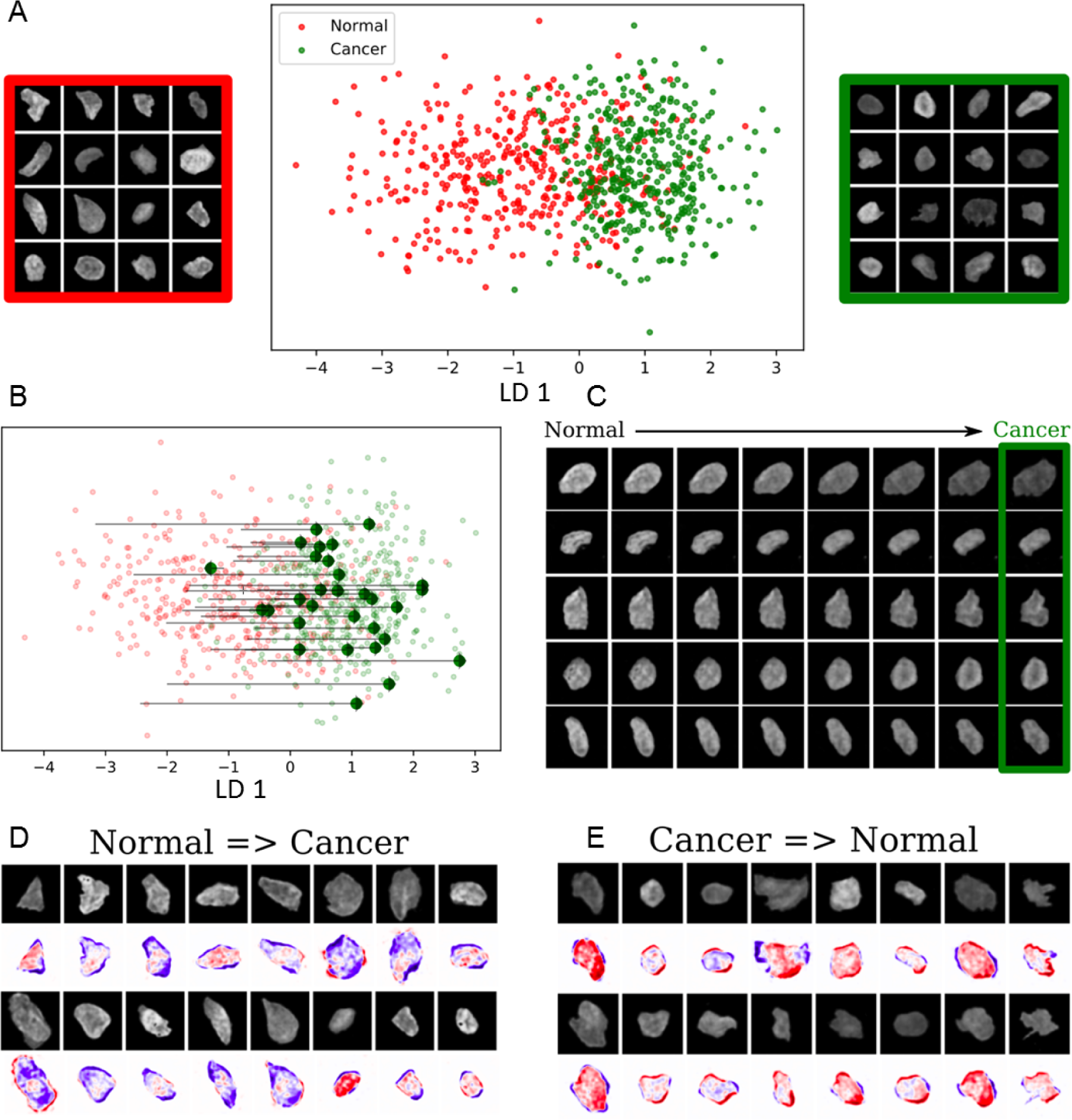
ImageAEOT applied to tracing cellular trajectories in breast tissues. (a) Visualization of normal and cancer nuclear images from tissue in both the original image space and the latent space using an LDA plot. (b) Predicted trajectories in the latent space using optimal transport. ImageAEOT was used to trace the trajectories from normal to cancer nuclei. (c) Predicted trajectories mapped back to the image space. Note that only the last image in each sequence is a real nucleus; the remaining images are predicted and generated by ImageAEOT. (d-e) Illustration of the principal features that change between normal and cancer nuclei, namely a combination of nuclear morphological and chromatin condensation features.

Understanding cell-state transitions during development or disease progression is a major challenge in modern biology and requires information on single-cell trajectories (21-23). However, given the long transition time scales for cell state switches, current experimental methods (including imaging, genomics and proteomics) are often limited to taking snapshots of different cells at different time points, making it a challenge to infer single-cell trajectories from such data. To address this challenge, we introduced ImageAEOT, which combines a variational autoencoder to map images to a latent space with optimal transport to model cell trajectories. While previous methods have used optimal transport for the analysis of single-cell RNA-seq data during differentiation (24), the use of a variational autoencoder is crucial when working with images to define a common coordinate system of the data (12).

Recent studies have revealed a strong coupling between cellular mechanical state, chromatin packing and the biochemical interaction networks within a cell (25-30). We therefore used nuclear images as a functional read-out to analyze cell-state transitions within the heterogeneous tissue microenvironment. In particular, we applied ImageAEOT to analyze the process of fibroblast activation and cancer progression, demonstrating that ImageAEOT can be used to identify the most salient image features of a particular cell state change without having access to continuous time data. Collectively, our results provide a quantitative framework to analyze cellular transitions at time scales relevant for developmental and disease processes, thereby opening novel routes for the identification of early disease biomarkers from imaging data.

## Acknowledgements

The authors thank Melanie Lee and Diego Pitta de Araujo for the schematic drawings. K.D.Y. was partially supported by the National Science Foundation (NSF) Graduate Research Fellowship and ONR (N00014-18-1-2765). The G.V.S. laboratory thanks the Mechanobiology Institute (MBI), National University of Singapore (NUS), Singapore and the Ministry of Education (MOE) Tier-3 Grant Program for funding. C.U. was partially supported by NSF (DMS-1651995), ONR (N00014-17-1-2147), and a Sloan Fellowship.

## Supplementary Materials

### Materials and Methods

#### ImageAEOT: Modeling Trajectories

Let 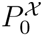 and 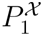 denote the distributions of cell images from state 0 and state 1 respectively, with the same support over the image space 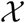. From here on, we refer to these as the source and target distributions. Predicting and modeling cell trajectories of cells from state 0 as they progress to state 1 can be formulated as learning a function 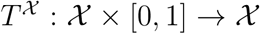 such that for 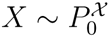, the following modeling conditions hold:

1. 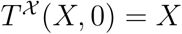
2. 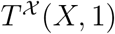 is semantically similar to both *X* and samples from 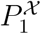
3. 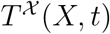 for *t* ∈ (0, 1) represents an intermediate transformation between 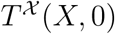 and 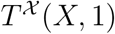.

Concretely, 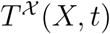, *t* ∈ [0, 1] models the trajectory of cell 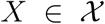 as it transitions from state 0 to state 1. For example, 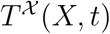, *t* ∈ (0, 1) could be the “shortest path” from 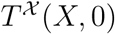 to 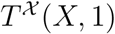. But since 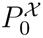 and 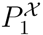 are supported on a low-dimensional manifold of 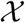, the shortest path between them in terms of Euclidean distance is not meaningful. Therefore, the first step of the ImageAEOT framework involves learning a pair of mappings that encode and decode 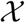 to a low-dimensional latent space 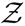, in which Euclidean distances between encodings reflect semantic distances between images. After the images have been embedded into such a latent space, ImageAEOT learns a transport map 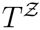 that pushes 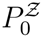 to 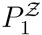 (i.e., the corresponding distributions in the latent space) in a way that minimizes the transport cost with respect to Euclidean distance. In summary, ImageAEOT completes the following three steps:

1. embedding cell images into a low dimensional latent space using a learned encoder *E*;
2. predicting how cells transform in the latent space by learning a transport map 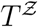; and
3. decoding cell embeddings back to images using a learned decoder *D*.

Composing these three maps yields a model of cell trajectories in the original image space, 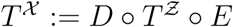 satisfying the modeling conditions above.

#### Embedding images into the latent space

To learn an encoder-decoder pair from the image space 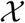 to a low-dimensional latent space 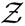 and back to the image space 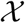, we use the following variational autoencoder (VAE) model (1). Suppose each image *X* is generated via the following process:

1. sample *Z* from *pz*(·), a prior distribution supported over 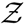;
2. sample *X* from *px*|*z*(·|*Z*), a distribution over 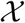 conditioned on *Z*.

In the VAE framework, *px*|*z*(·|*Z*) is modeled by the conditional distribution 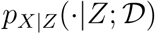, which is parametrized by the *decoder neural network* 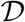. Therefore, an image *X* can be generated from a latent encoding *Z* by sampling 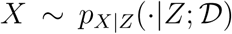. We define this as our probabilistic decoder map *D*. Additionally, the VAE approximates the posterior distribution 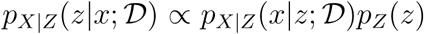 by the variational distribution 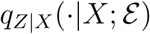, which is parametrized by the *encoder neural network* 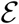. To encode an image *X* to the latent space, we sample 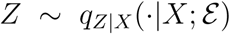. We define this as our probabilistic encoder map *E*. The networks 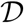 and 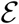 are optimized by maximizing a lower bound on the probability of observing from the empirical distribution of training images 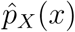 under the generating distribution 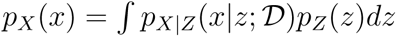. This lower bound is given by
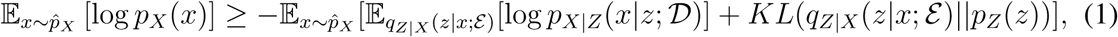

where *KL* is the Kullback-Leibler divergence. To simplify computations, 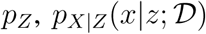, and 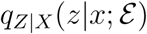 are modeled as multivariate Gaussians with diagonal covariance matrices, and then 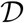, 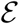 are optimized by doing stochastic gradient descent on (1). In practice, poor reconstruction quality can result from directly optimizing (1), and this can be ameliorated by tuning a hyperparameter *λ* ∈ (0, 1] before the KL-divergence term.

#### Learning a transport map in the latent space

Once the source and target images are embedded into the latent space using the variational autoencoder from the previous section, ImageAEOT learns a transport map 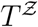 from the source distribution to the target distribution in the latent space using the principles of optimal transport. Importantly, this mapping is optimal with respect to the Euclidean distance in the latent space and can be computed as follows.

Formally, let 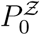 and 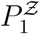 be the distributions of images embedded in the latent space, and let 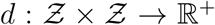 be the Euclidean distance. First we solve the Kantorovich optimal transport problem: find a probabilistic coupling *γ*^∗^ (i.e. a joint distribution over 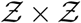) whose marginals are 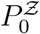 and 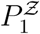 that minimizes

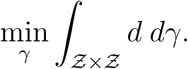

When 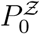 and 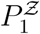 are discrete, an approximation of *γ*^∗^ can be computed efficiently using the Sinkhorn-Knopp algorithm (2), shown in Algorithm 1. The inputs are *r*, *c*: vectors summing to 1 representing the probability mass functions over the supports of 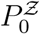 and 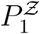 respectively; and *C*: a distance matrix whose *ij*-th entry equals the distance between the *i*-th point in the support of 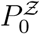 and the *j*-th point in the support of 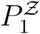. The output is a matrix *γ*^∗^ representing a probabilistic coupling between 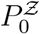 and 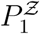. Once *γ*^∗^ is obtained, for each *Z* sampled from 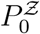, 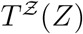 is computed by taking the mean of the support of 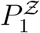 weighted based on *γ*^∗^. Specifically, for the *i*-th point in 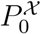 denoted by *Z_i_*, we have that

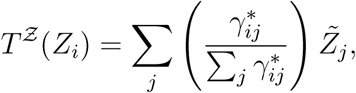

where 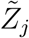 are the points in 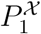.

##### Algorithm 1 Sinkhorn-Knopp algorithm

**INPUT:** Probability mass functions *r*, *c*, distance matrix *C*, tuning paramter *λ*

**OUTPUT:** Probabilistic coupling *γ*^∗^

1: Compute *K* := exp [−*λC*]

2: Initialize *u*, *v*

3: **while** not converged **do**

4:         Update *v* = *c*./*K^T^u*

5:         Update *u* = *r.*/*Kv*

6: **end while**

7: Return *γ*^∗^ := *diag*(*u*)*Kdiag*(*v*)

### ImageAEOT: Identifying Biomarkers via Perturbations

To identify biomarkers of the target population 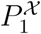, ImageAEOT takes a sample from the source population 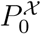 and endows it with the characteristics of 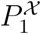. This is done by encoding the sample to the latent space, performing a small perturbation in the direction of 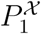, and then decoding to the image space. Concretely, the function *T_p_* that maps each sample to a perturbed sample in the latent space is defined as
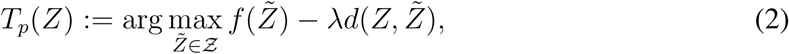

where *d* is the (Euclidean) distance function, *λ* > 0 is a parameter that reflects the size of the perturbation, and *f*(·) is a function that maps points in 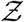 to how similar they are to samples from 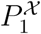. In practice, we either chose *f*(·) to be (i) the first discriminant function from a linear discriminant analysis model of 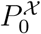 and 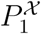; or (ii) the log-probability that *Z* is drawn from state 1 based on a neural network classifier trained on 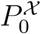 and 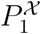.

### Implementation

For performing latent discriminant analysis in our pipeline, we used the Python scikit-learn library (3). Neural network models were implemented in Python using the Pytorch library (4) and trained on an NVIDIA GeForce GTX 1080TI graphics card. The architecture of the VAE model in the ImageAEOT is shown in Supplementary Figure 3. Hyperparameter values of 0, 1e-6, 1e-7, 1e-8 were used in the objective. This model was trained using the Adam optimizer, a popular variant of stochastic gradient descent (5), with learning rate initialized at 1e-4, and batch sizes of 64 images. For classification tasks in the latent space, we implemented either a linear network or a feedforward network with ReLU activations and two hidden layers. The models were trained using the Adam optimizer with learning rate initialized at 1e-3. The VoxNet model architecture (6) used for classifying whole images is shown in Supplementary Figure 7. For the feature ablation studies, feature extraction was done using the Python mahotas library (7), and the logistic regression model was implemented using scikit-learn (3).

## Cell-Culture Experiments

### Reagents used

NIH3T3 (CRL-1658), BJ (CRL-2522), MCF10A (CRL-10317), MCF7 (HTB22) and MDA-MB-231 (HTB-26) cells were obtained from ATCC. They were cultured in DMEM-high glucose (ThermoFisher Scientific 11965092) media supplemented with 10%FBS (ThermoFisher Scientific 16000044) and 1% pen-strep (Sigma P4333) antibiotic. Antibodies used: EpCAM (Cell Signaling Technology, 2929S) and Vimentin (Cell Signaling Technology, 5741S). Other reagents: Breast tissue sections within normal limits (CS708873, Origene), metastatic breast adenocarcinoma tissue sections (CS548359, Origene), Histozyme (H3292-15ML, Sigma), Pro-Long Gold Antifade Mountant with DAPI (P36941, ThermoFischer Scientific), Paraformaldehyde PFA (Sigma, 252549-500ml), Triton (Sigma, X100-100ml) and DAPI solution (ThermoFisher Scientific R37606) and IF wash buffer (for 250ml: 125mg NaN3 + 500l Triton X-100 + 500l Tween-20 in 1X PBS).

### Micro-contact printing

Fibronectin micropatterning was performed as described in (8). Briefly, circular fibronectin (Sigma F1141-2MG) micropatterns (area = 1800m2) were made on uncoated Ibidi dishes (81151). These micropatterned dishes were then passivated with 0.2% pluronic acid (Sigma P2443) for 10 minutes and washed with PBS.

### Co-culture experiment

Cell culture and experiments were all performed at 37C, 5% CO2 and humid conditions. MCF7 cells were seeded previous day (Day −1) on 1800 m2 circular fibronectin micropatterns. After 24 hours, MCF7 clusters (cluster of 10 cells) were obtained. Collagen gel (1mg/ml) was prepared as per the manufacturers instructions along with DMEM media. MCF7 clusters were then scraped off and mixed with 30,000 NIH3T3 fibroblasts in 300l of collagen gel solution which was then added to an uncoated Ibidi dish. For control conditions, either only MCF7 clusters or 30,000 NIH3T3 cells were added with 300l of collagen gel solution in each Ibidi dish. The gel was then allowed to solidify at 37C. After two hours, 500l of fresh media was added to these samples. Each condition was prepared as shown in Table 1. Each set of dishes were then fixed on Day1, Day2, Day3 and Day4.

**Table 1:** Description of co-culture experiment.

| Conditions | Control NIH3T3 | Control MCF7 | Co-culture (NIH3T3 and MCF7) |
| --- | --- | --- | --- |
| Day -1 | | MCF7 clusters seeded on $1800\mu m^2$ circles | MCF7 clusters seeded on $1800\mu m^2$ circles |
| Day 0 | 30,000 NIH3T3 cells added | MCF7 clusters of about 10 cells from each micropattern mixed with collagen | MCF7 clusters of about 10 cells from each micropattern mixed with collagen and 30,000 NIH3T3 cells added |
| Days 1-4 | Cells fixed | Cells fixed | Cells fixed |

## IF Staining

For IF staining, media was aspirated and 4%PFA was added and incubated for 20 minutes. The samples were then washed thrice with PBS + glycine buffer. The gel was treated with 0.5% Triton for 20 minutes to permeabilize the cells followed by washes with the PBS + glycine buffer. The samples were then blocked with 10% goat serum in IF wash buffer (blocking solution) for 2 hours. Primary antibodies in blocking solution were then incubated overnight as per the dilution recommended by the manufacturer. Next day, the samples were washed thrice with IF wash buffer for 5 minutes each. Secondary antibody in blocking solution was then added as per the manufacturers instruction for 2 hours. Samples were then washed thrice with IF wash buffer for 5 minutes each. DAPI solution was added to the samples and stored temporarily at 4C until imaged.

## Immunohistochemistry

Formalin-fixed, paraffin-embedded (FFPE) tissue sections (5m thickness) on slides were deparaffinized by heating them in an oven at 60C for 5 minutes and subsequently washing them with xylene. The sections were then rehydrated in serially diluted ethanol solutions (100% - 50%) as per standard protocols and rinsed with water. Antigen retrieval was performed using Histozyme solution as per the manufacturers protocol and then rinsed with water. DAPI was then added to these sections and they were covered with a coverslip. The slides were incubated for 24 hours after which the coverslips were sealed and taken for imaging.

## Imaging

Most of the solution in the dish was aspirated before imaging. Around 50l of the solution was left to prevent drying of the collagen gel. The images were obtained using a Nikon A1R confocal microscope. For co-culture samples, Z-stack images were captured using 40X objective (water, 1.25 NA), 0.3*µm* pixel, Z-depth of 1*µm* and all images were captured for not more than 50*µm* depth. Each image is 1024X1024 pixels in size. For larger field images, 2 × 2 images or 3 × 3 images were obtained and stitched together with 5% overlap. For tissue slices, wide-field images were obtained using an Applied Precision DeltaVision Core microscope with 100X objective (oil, NA 1.4) and a pixel size of 0.2150*µm*. These 512 × 512 12-bit images were deconvolved (enhanced ratio, 10 cycles) and saved in .tiff format.

## Segmentation of Nuclei

Images were analyzed using custom codes written in ImageJ2/Fiji (9). The raw 3D images labelled for DNA using DAPI, acquired using a laser scanning confocal microscope, were filtered using a Gaussian blur and thresholded using automated global thresholding method such as Moments to binarize and identify nuclear regions. Watershed was used to separate touching nuclear regions. This binary image was then used to identify individual nuclei as 3D objects within a size range of 200-1300*µm*^3^. Each nucleus identified as a separate 3D object was visualized with distinct colors. In order to smoothen any irregular boundaries, a 3D convex hull was carried out and then the individual nuclei were cropped along their bounding rectangles and stored. This was carried out using the functions from Bioformats and the mcib3d library. In order to separate nuclei that were clumped and could not be separated using watershed, the 3D Euclidian distance transform was carried out on these clumps of nuclei followed by a second round of thresholding to remove pixels from the boundaries; then individual nuclei were identified as described earlier. From this set, the blurred out of focus nuclei and the over-exposed nuclei were filtered out and then the selected nuclei were used for further analysis.

## Supplementary Figure Legends

**Supplementary Figure S1:**
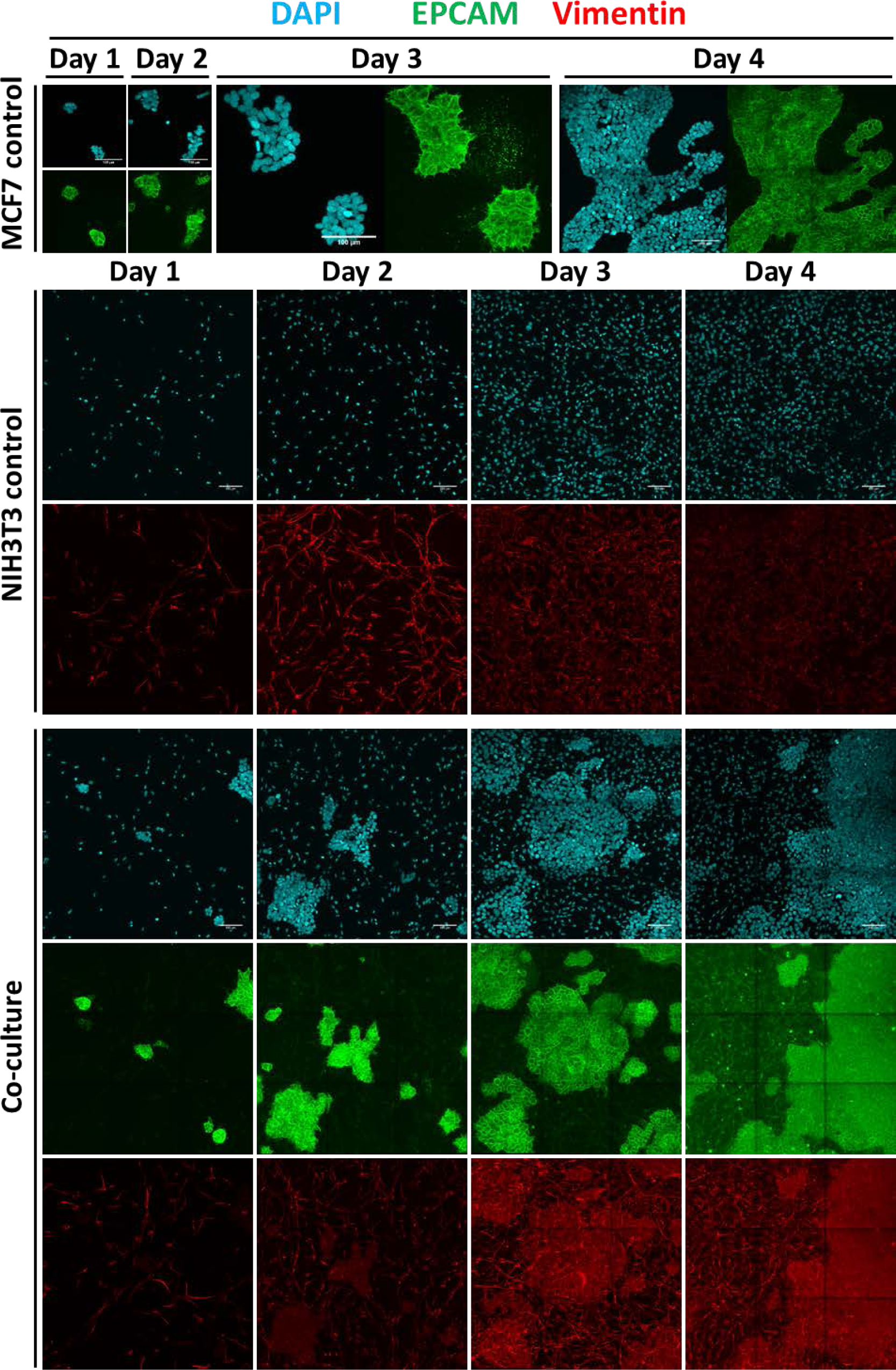
Co-culture experiment. Representative maximum intensity projected images of control MCF7 cells, control NIH3T3 cells and MCF7-NIH3T3 co-culture cells in 3D collagen gel from Day 1 to Day 4. EpCAM (green) is enriched in MCF7 cells, Vimentin (red) is enriched in NIH3T3 cells; the nuclei are also labelled using DAPI (blue). Each image acquired is 1024 x 1024 pixels in size. To obtain a larger field of view, multiple images were stitched together to create a large image, consisting of 4 images (2 x 2) for Day 3 and Day 4 in the control MCF7 condition and 9 images (3×3) in the control NIH3T3 and co-culture conditions. Scale bar: 100μm.

**Supplementary Figure S2:**
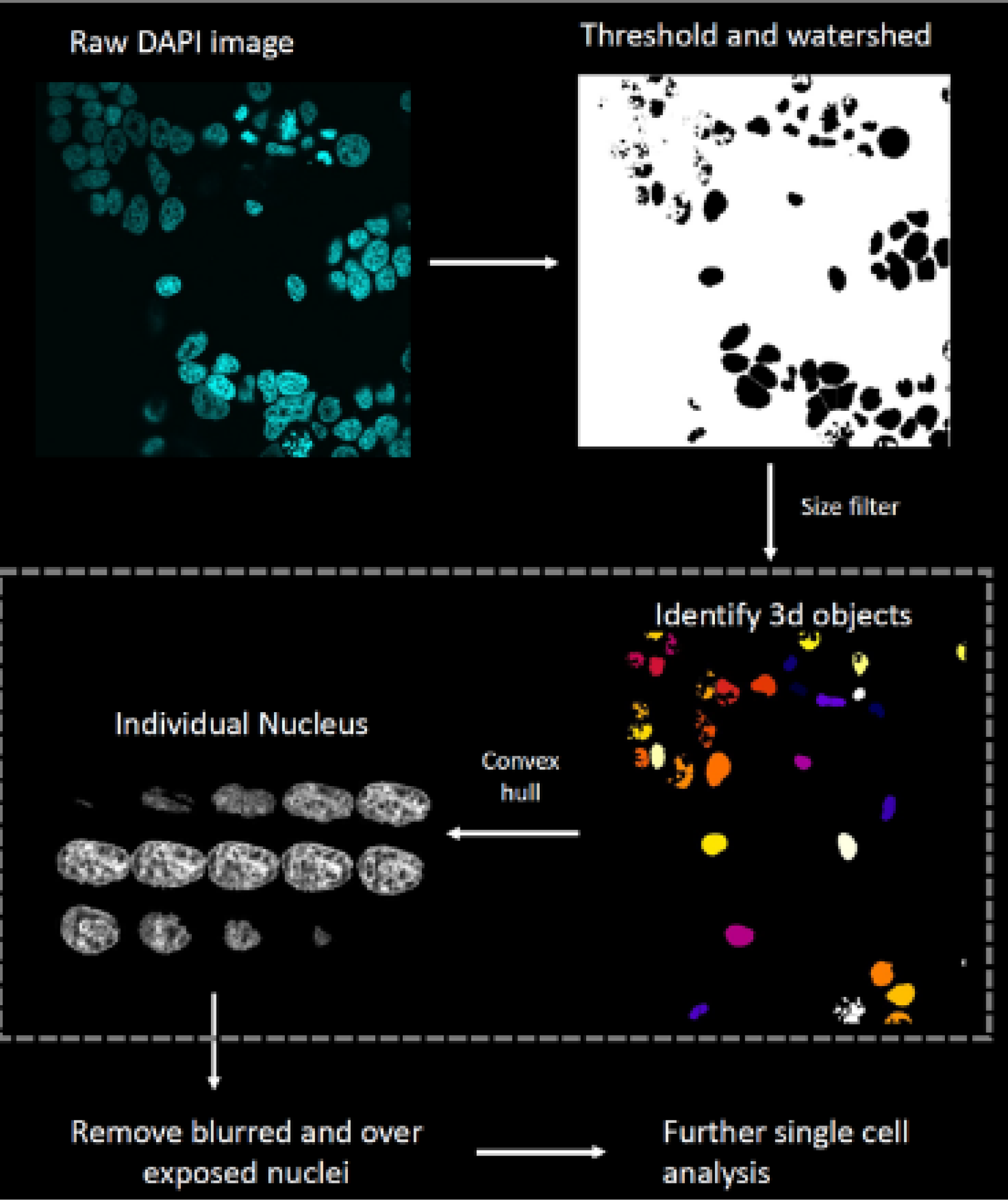
Image processing workflow: The raw 3D images labelled for DNA using DAPI, acquired using a laser scanning confocal microscope, are filtered using a Gaussian blur and thresholded using an automated global thresholding method such as otsu to binarize the image and identify nuclear regions. Watershed is used to separate closeby nuclei. The resulting binary image is then used to identify individual nuclei as a 3D objects within a size range of 200-1300μm^3^. Each nucleus identified as a separate 3D object is visualized with distinct colors. In order to smoothen any irregular boundaries, a 3D convex hull is constructed and then the individual nuclei are cropped along their bounding rectangles and stored. From this set, the blurred out of focus nuclei or over-exposed nuclei are filtered out and then the remaining nuclei are used for further analysis.

**Supplementary Figure S3:**
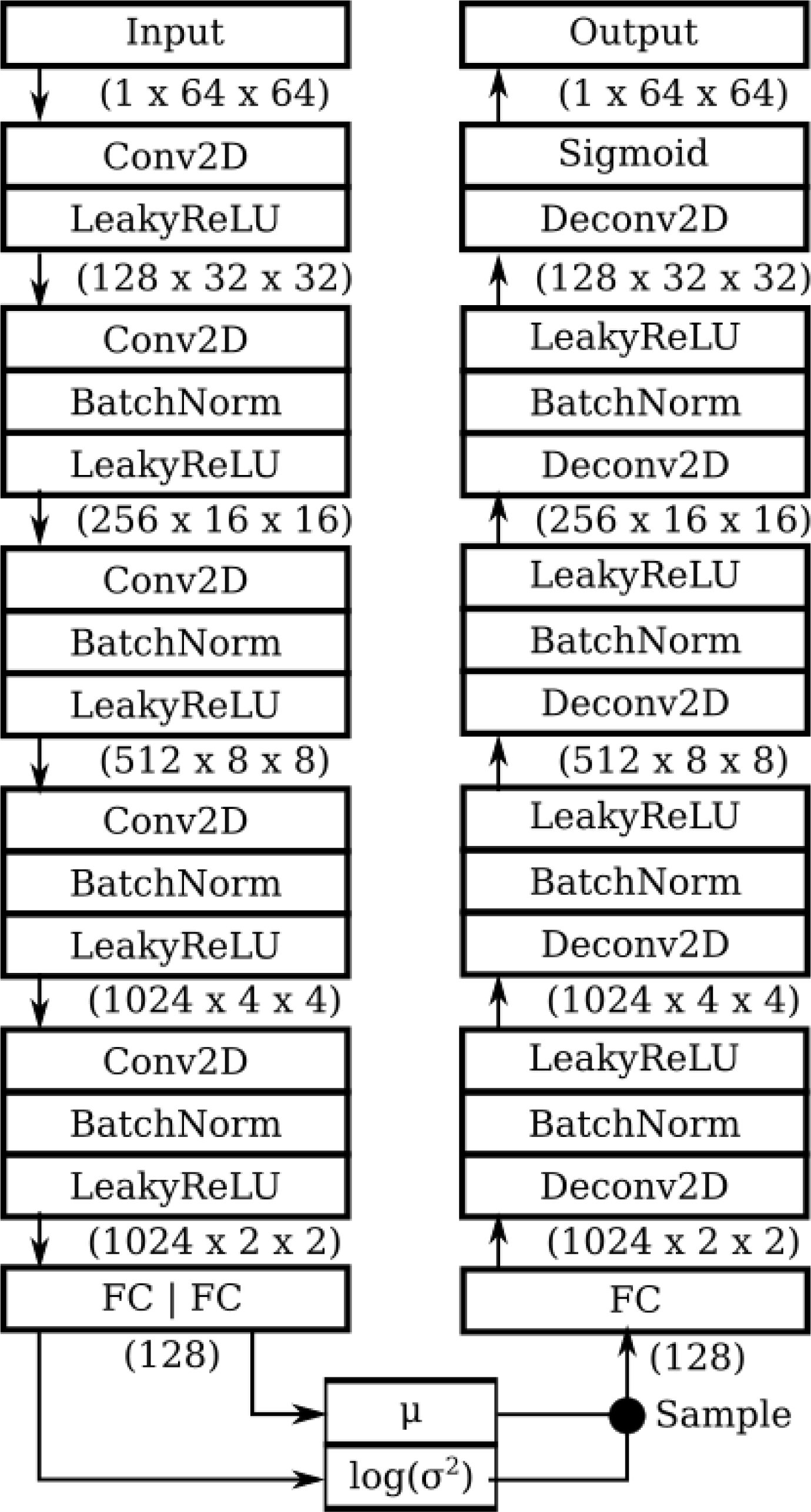
Architecture of variational autoencoder. The encoder used for mapping images to the latent space is shown on the left. This encoder takes images as input and returns Gaussian parameters in the latent space that correspond to this image. The decoder used for mapping from the latent space back into the image space is shown on the right.

**Supplementary Figure S4:**
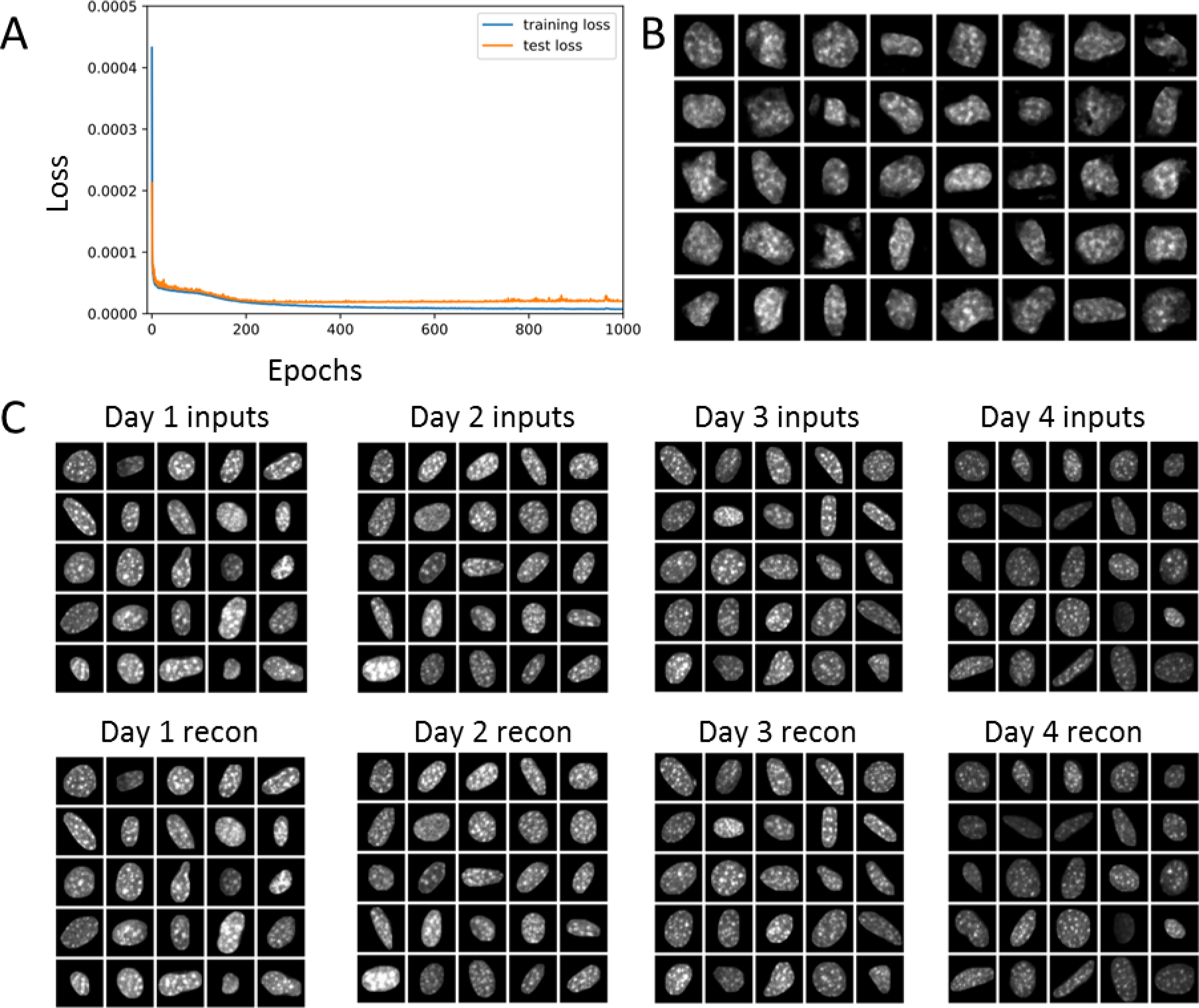
Training the variational autoencoder on co-culture NIH3T3 nuclei. (a) Training and test loss curves of the variational autoencoder plotted over 1000 epochs. (b) Nuclear images generated from sampling random vectors in the latent space and mapping these to the image space. These random samples resemble nuclei, suggesting that the variational autoencoder learns the manifold of the image data. (c) Input and reconstructed images from Day 1 to Day 4 illustrating that the latent space captures the main visual features of the original images.

**Supplementary Figure S5:**
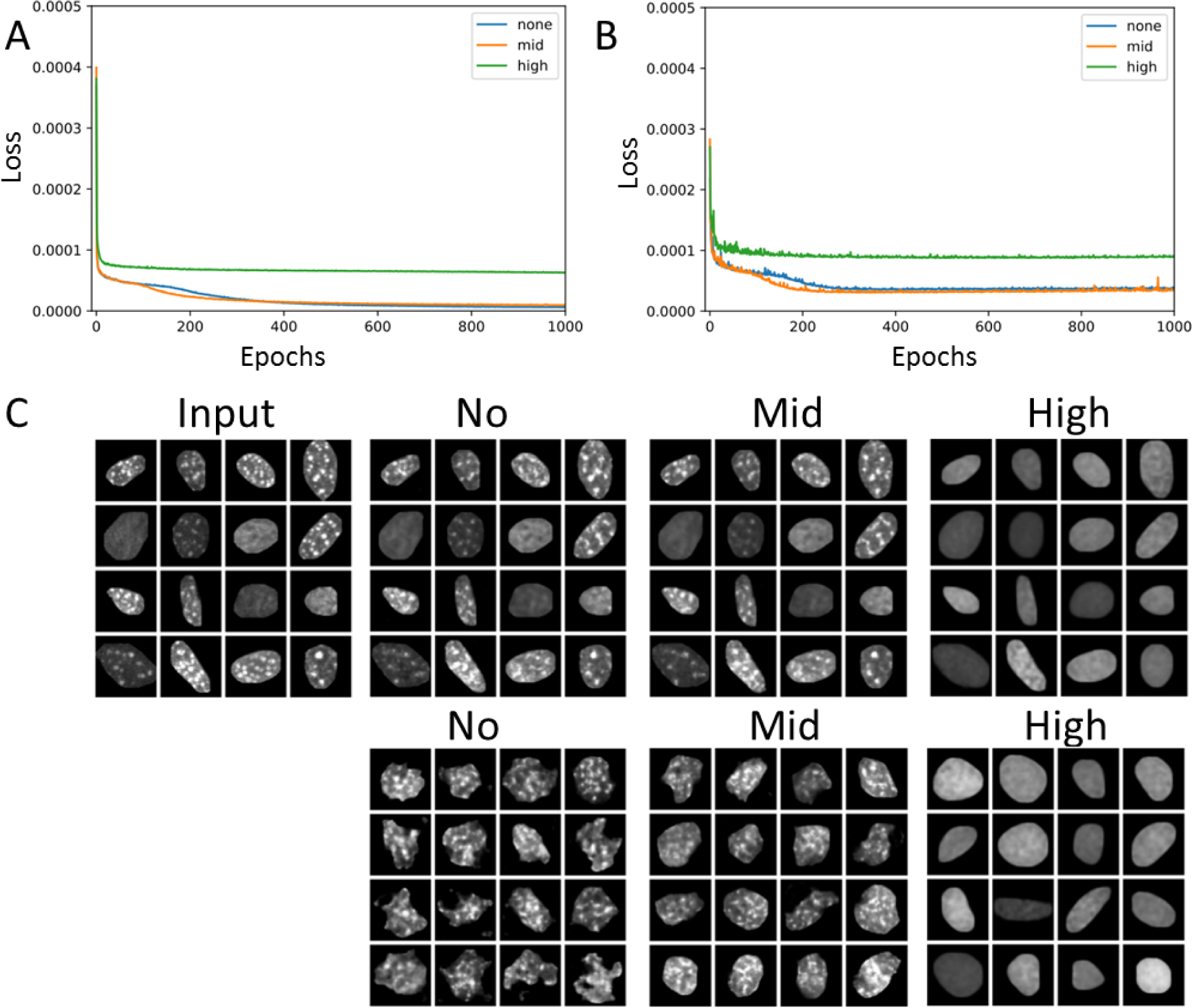
Hyperparameter tuning for the variational autoencoder over co-culture nuclei. (a-b) Training loss and test loss curves respectively for high, mid, or no regularization. (c, top row) Reconstruction results for each model. Models with no or mid-level regularization can reconstruct input images well, while models with high regularization do not. (c, bottom row) Sampling results for each model. Models with no regularization do not generate random samples as well as models with mid-level regularization, which suggests that the model with mid-level regularization best captures the manifold of nuclei images.

**Supplementary Figure S6:**
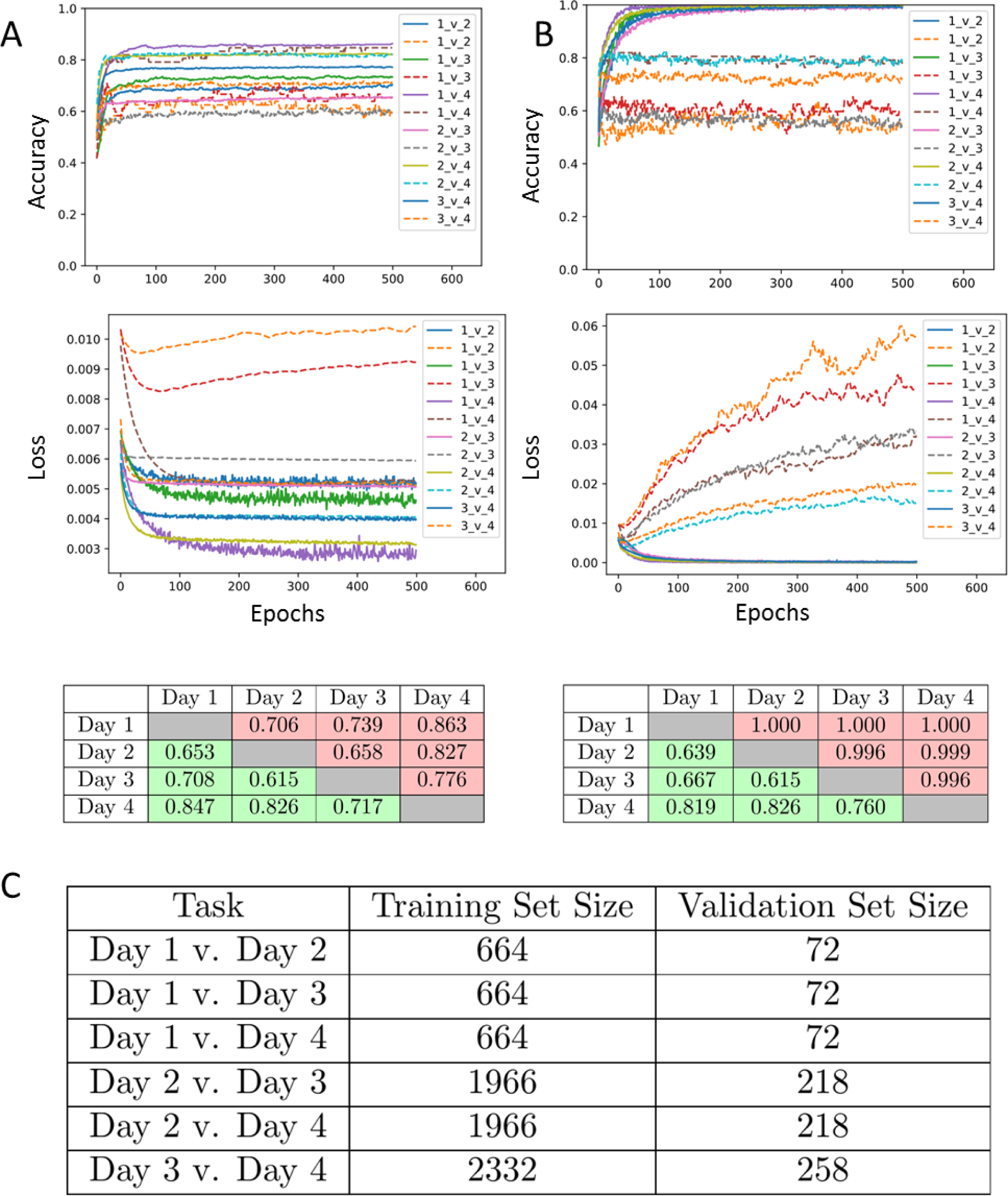
Pairwise classification of NIH3T3 cells from co-culture model in the latent space. (a) Classification results in the latent space using a linear model. Top: training and test loss curves for each pairwise comparison (Day 1, Day 2, Day 3, Day 4). Middle: training and test accuracy curves for each pairwise comparison. Bottom: Table of best training (red) and test (green) accuracy for each classification task. (b) Same as (a) but using a two-layer feedforward neural network. (c) Training and validation dataset sizes for each of the classification tasks.

**Supplementary Figure S7:**
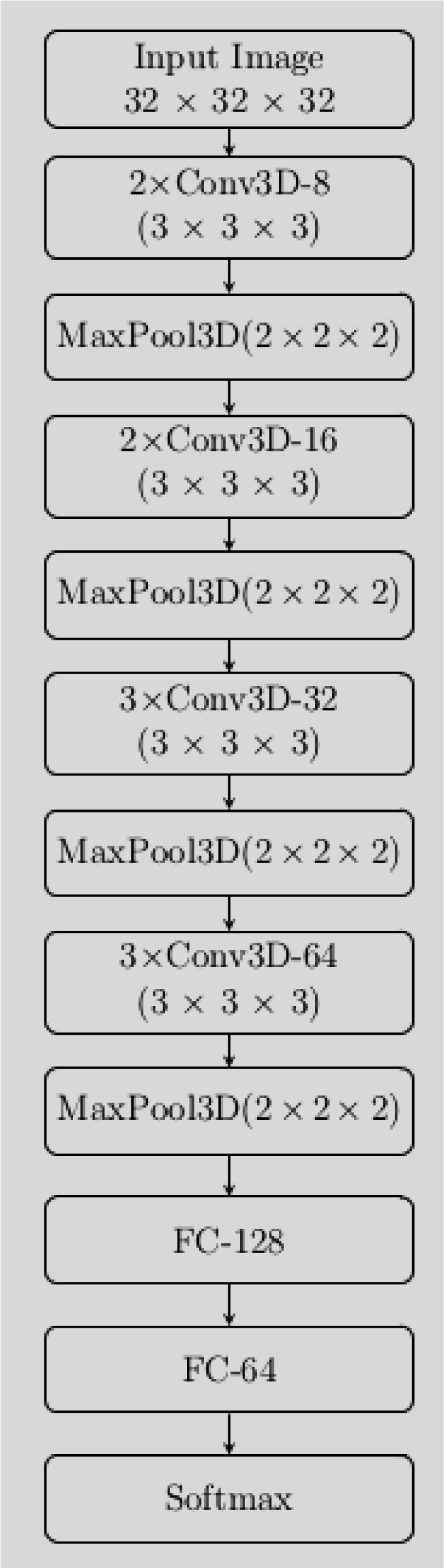
VoxNet architecture used in the classification tasks. The input images are of size 32 × 32 × 32. The notation *r*×Conv3D-*k* (3 × 3 × 3) means that there are *r* 3D convolutional layers (one feeds into the other) each with *k* filters of size 3×3×3. MaxPool3D(2×2×2) indicates a 3D max pooling layer with pooling size 2 × 2 × 2. FC-k indicates a fully connected layer with k neurons. Note that the PReLU activation function is used in every convolutional layer while ReLU activation functions are used in the fully connected layers. Finally, batch normalization is followed by every convolutional layer.

**Supplementary Figure S8:**
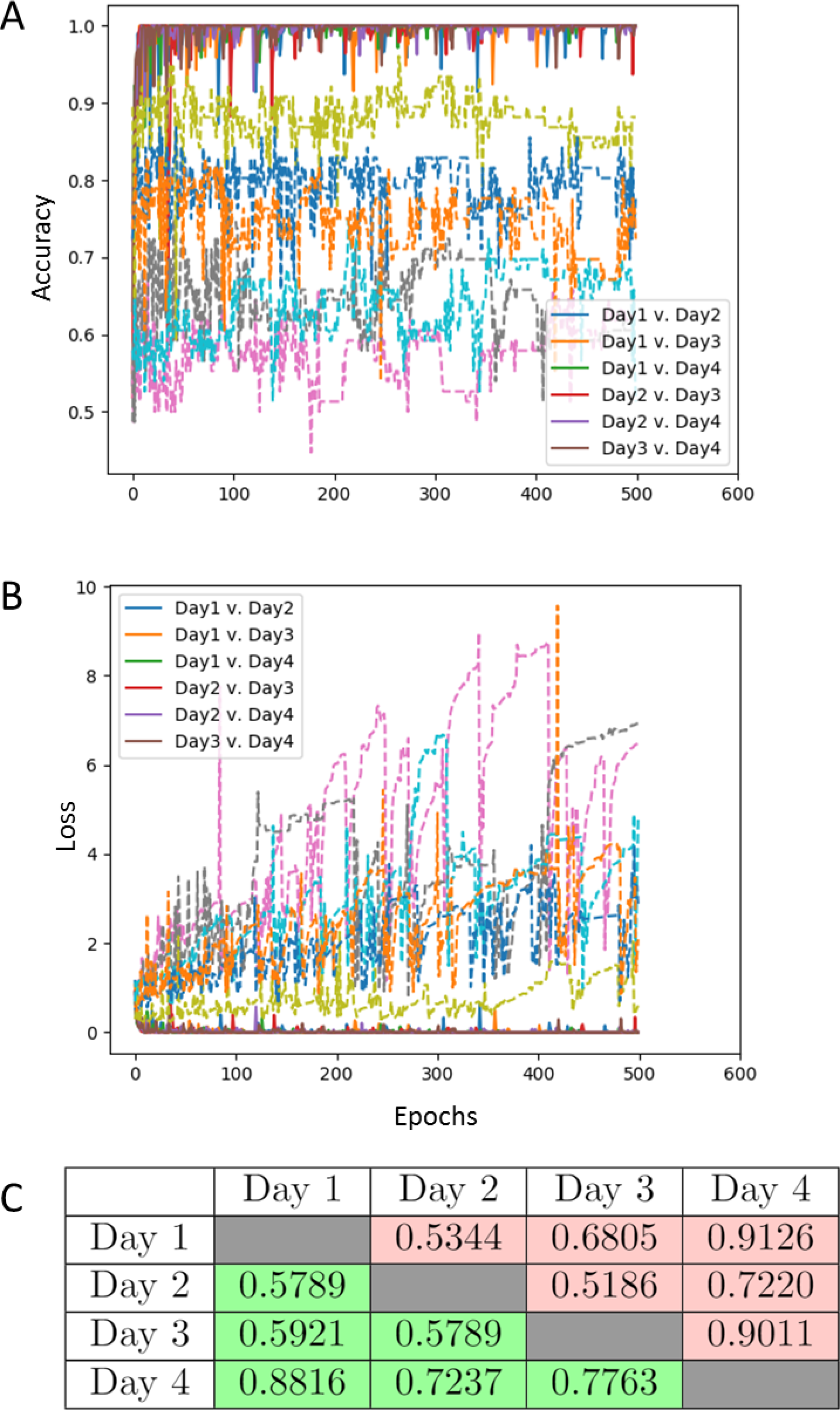
Pairwise classification of NIH3T3 cells from co-culture model using VoxNet. (a) Accuracy and (b) loss curves for each pairwise comparison (Day 1, Day 2, Day 3, Day 4). (c) Table of best training (red) and test (green) accuracy for each classification task.

**Supplementary Figure S9:**
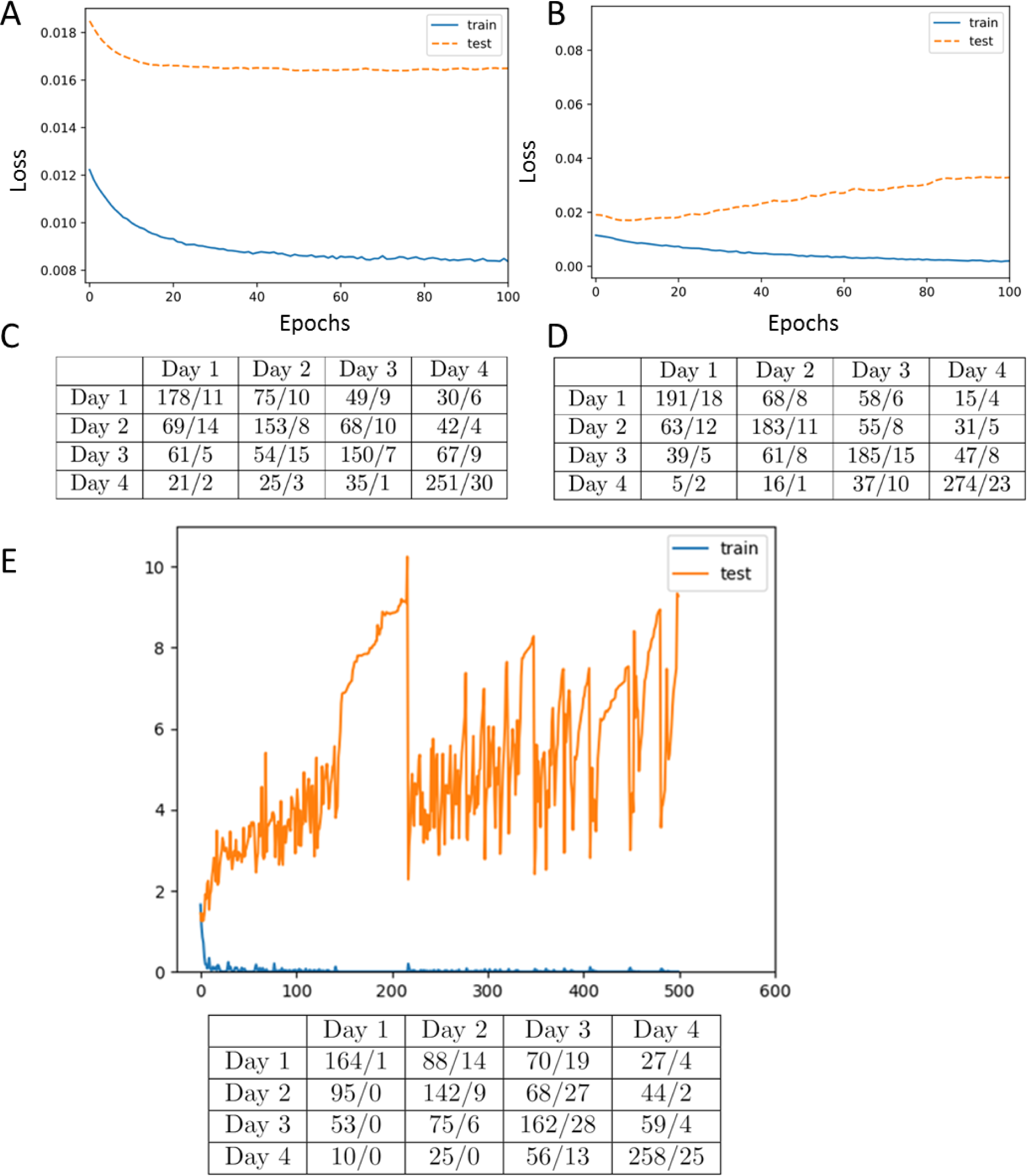
4-way classification results for co-culture NIH3T3 nuclei. Training and test loss curves for 4-way classification task (Day 1, Day 2, Day 3, Day 4) of co-culture NIH3T3 nuclei in the latent space using a linear model (a) and 2-layer feedforward neural network (b). (c-d) Confusion matrices for the classification tasks in (a-b). Each entry (X/Y) in row “A” and column “B” indicates that X nuclei of class “A” were classified as “B” in the training set and Y nuclei of class “A” were classified as “B” in the test set. (e) Same as (b,d) but for the 4-way classification task in the original image space using a deep convolutional neural network.

**Supplementary Figure S10:**
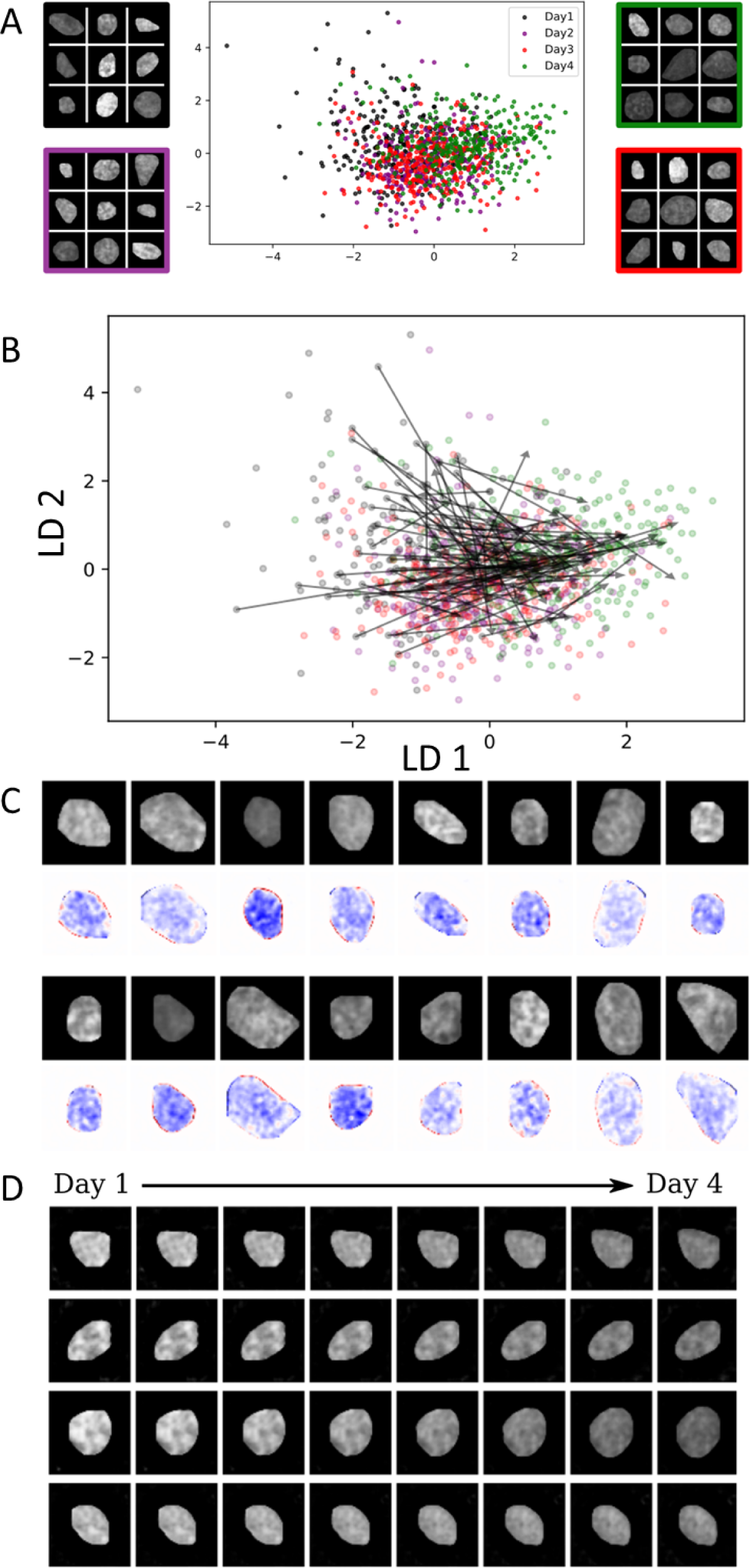
ImageAEOT applied to tracing trajectories of cancer cells in a co-culture system. (a) Visualization of MCF7 nuclear images from Days 1-4 in both the image and latent space using an LDA plot. Note that the distributions of the data points in the LDA plot appear to coincide, suggesting that the MCF7 cells do not undergo drastic changes from Day 1 to 4. Day 1: black; Day 2: purple; Day 3: red; Day 4: green. (b) Predicted trajectories in the latent space using optimal transport. ImageAEOT was used to trace the trajectories of Day 1 MCF7 to Day 4 MCF7. Each black arrow is an example of a trajectory. (c) Visualization of the principal feature along the first linear discriminant. The nuclear images are of Day 1 MCF7 cells. The images below show the difference between the generated images along the first linear discriminant and the original image (blue: decrease in pixel intensity; red: increase in pixel intensity). These results suggest that MCF7 nuclei do not exhibit drastic changes other than a reduction of intensity. (d) Predicted trajectories mapped back to the image space. Note that only the first image in each sequence is a real Day 1 MCF7 nucleus; the remaining images are predicted and generated by ImageAEOT. Note that there are only small changes in the nuclei, other than a decrease in overall intensity.

**Supplementary Figure S11:**
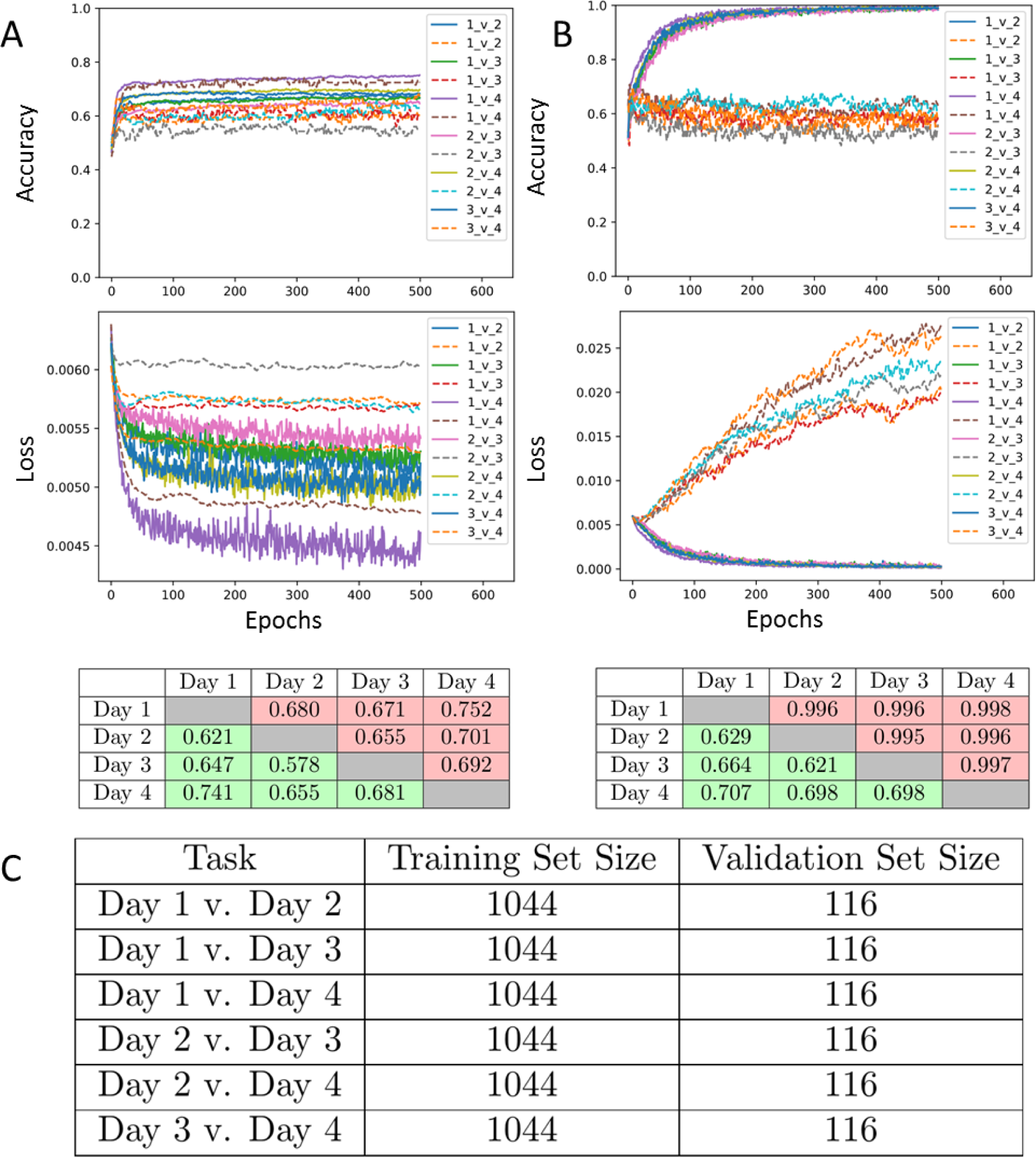
Pairwise classification of MCF7 cells from co-culture model. (a) Classification results in the latent space using a linear model. Top: training and test loss curves for each pairwise comparison (Day 1, Day 2, Day 3, Day 4). Middle: training and test accuracy curves for each pairwise comparison. Bottom: Table of best training (red) and test (green) accuracy for each classification task. (b) Same as (a) but using a two-layer feedforward neural network. (c) Training and validation dataset sizes for each of the classification tasks.

**Supplementary Figure S12:**
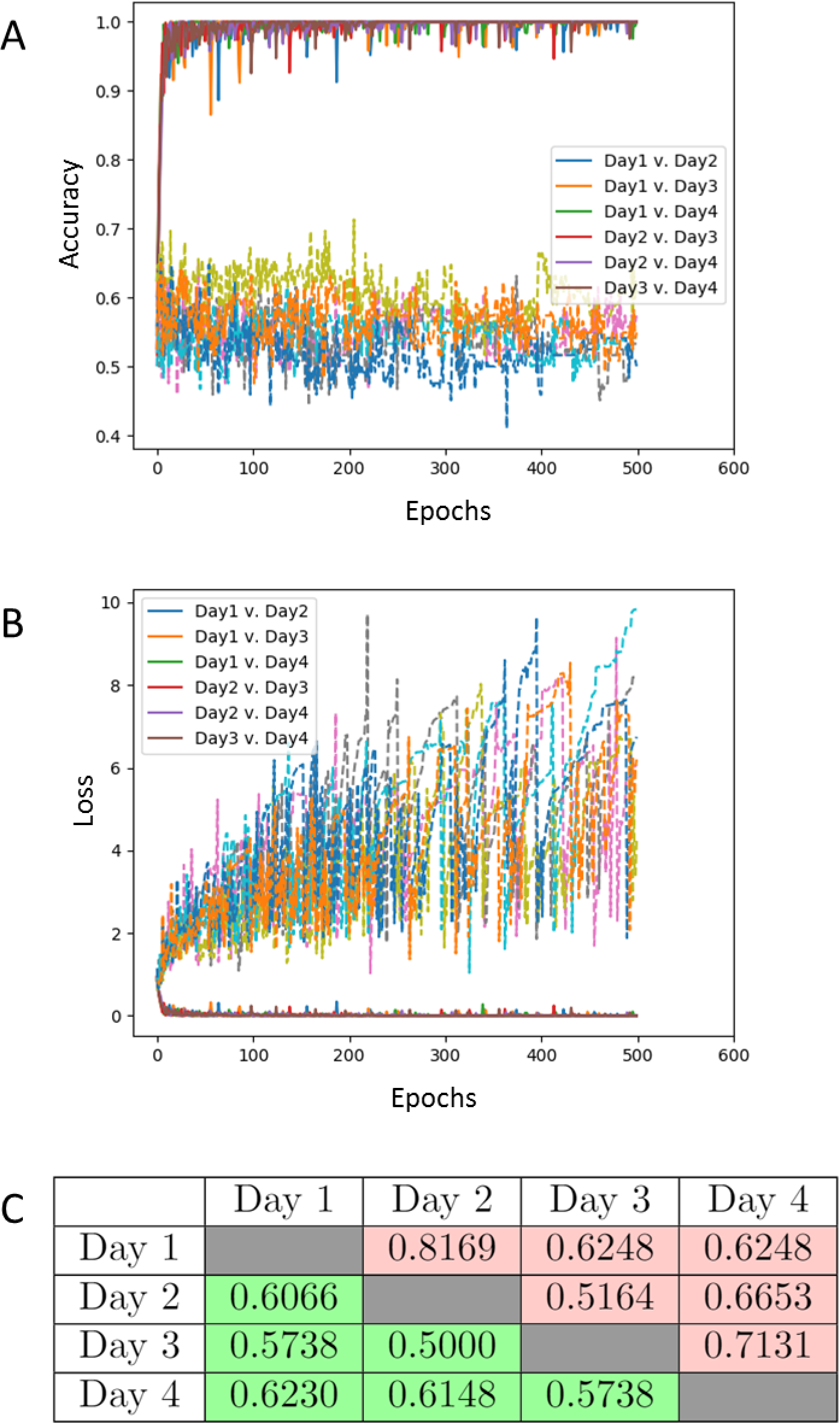
Pairwise classification of MCF7 cells from co-culture model using VoxNet. (a) Accuracy and (b) loss curves for each pairwise comparison (Day 1, Day 2, Day 3, Day 4). (c) Table of best training (red) and test (green) accuracy for each classification task.

**Supplementary Figure S13:**
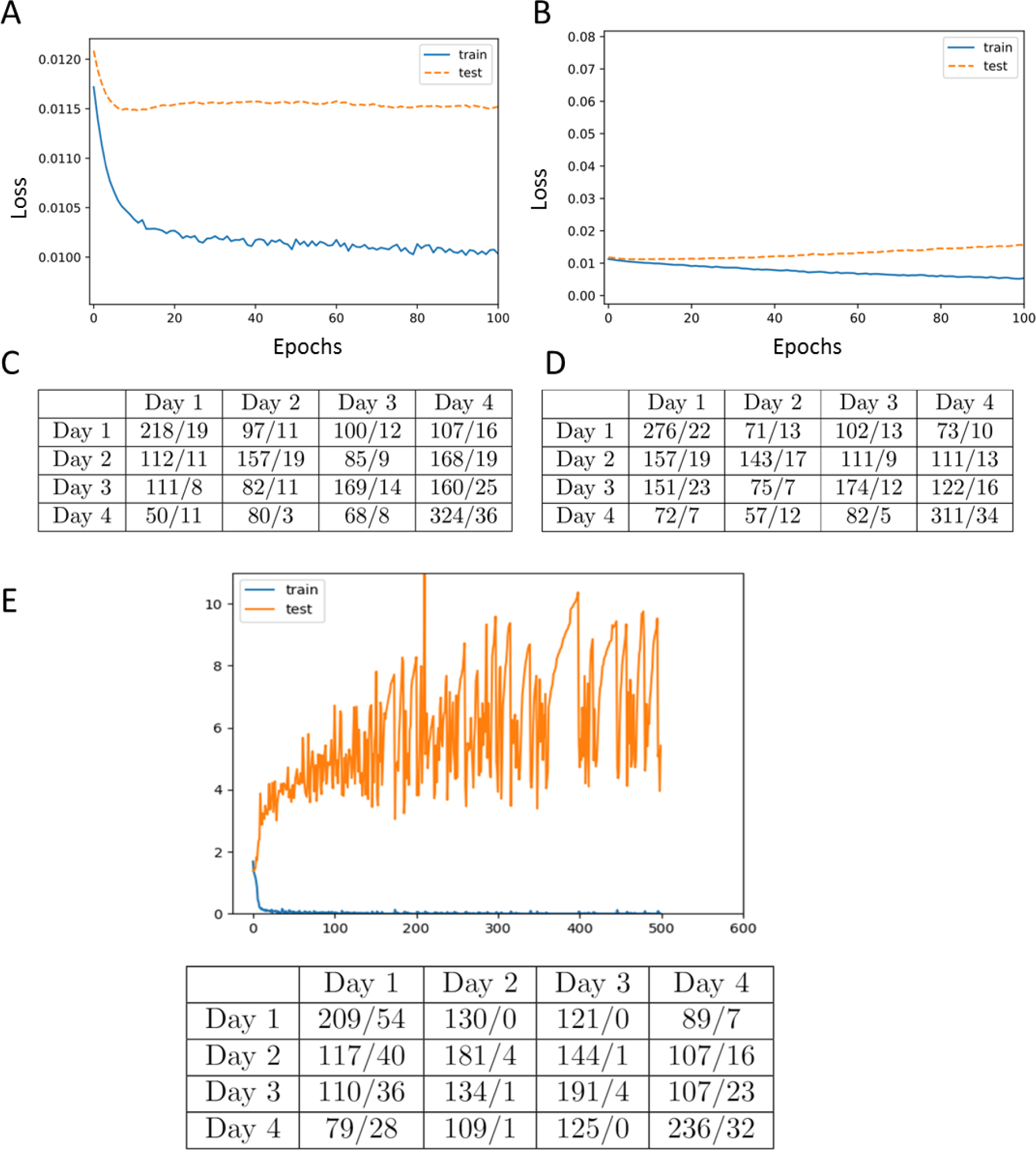
4-way classification results for co-culture MCF7 nuclei. Training and test loss curves for 4-way classification task (Day 1, Day 2, Day 3, Day 4) of co-culture MCF7 nuclei using a linear model (a) and 2-layer feedforward neural network (b). (c-d) Confusion matrices for the classification tasks in (a-b). Each entry (X/Y) in row “A” and column “B” indicates that X nuclei of class “A” were classified as “B” in the training set and Y nuclei of class “A” were classified as “B” in the test set. (e) Same as (b,d) but for the 4-way classification task in the original image space using a deep convolutional neural network.

**Supplementary Figure S14:**
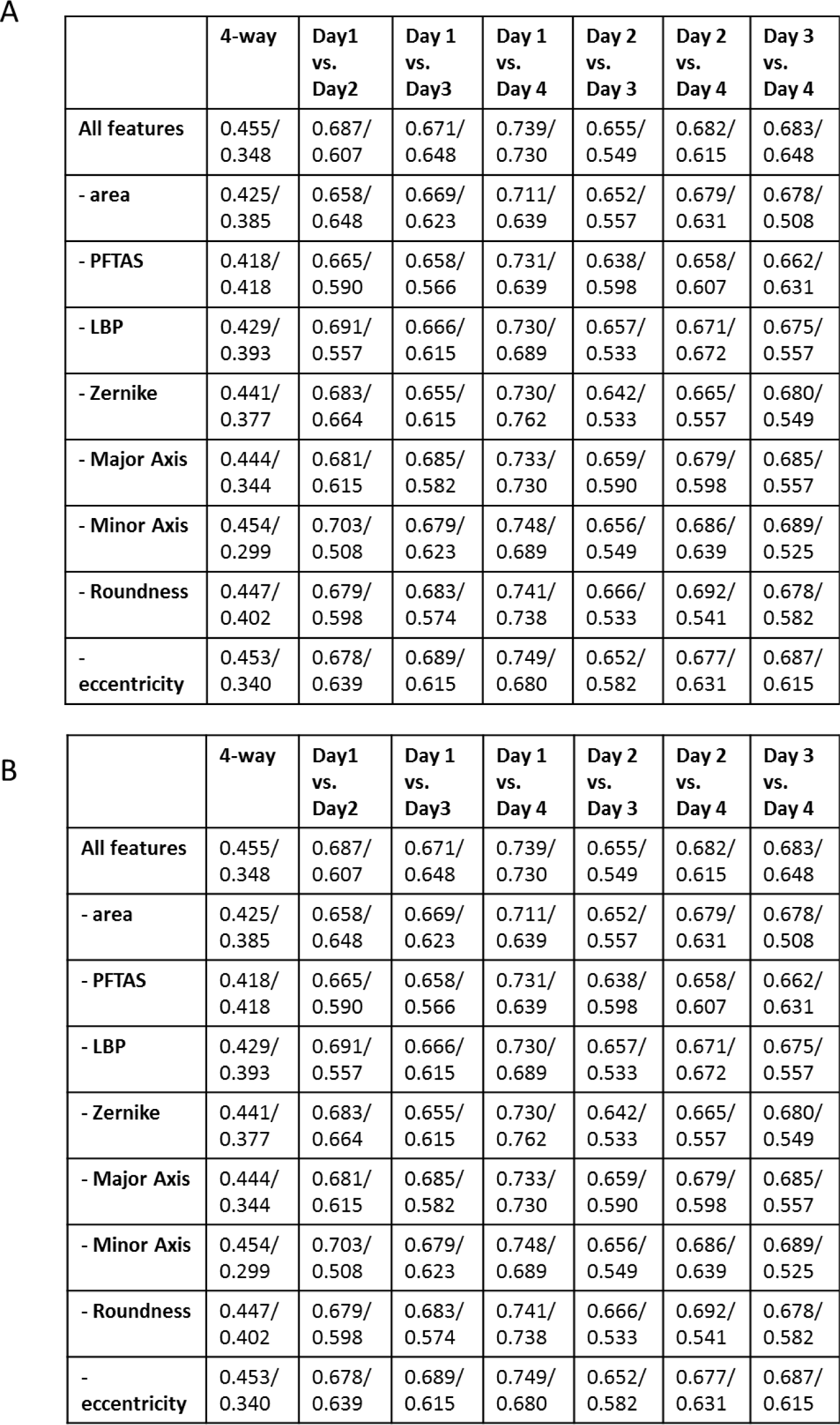
Feature ablation tables for co-culture nuclei. (a) Feature ablation table for logistic regression on NIH3T3 co-cultured cells. (b) Feature ablation table for logistic regression on MCF7 co-cultured cells.

**Supplementary Figure S15:**
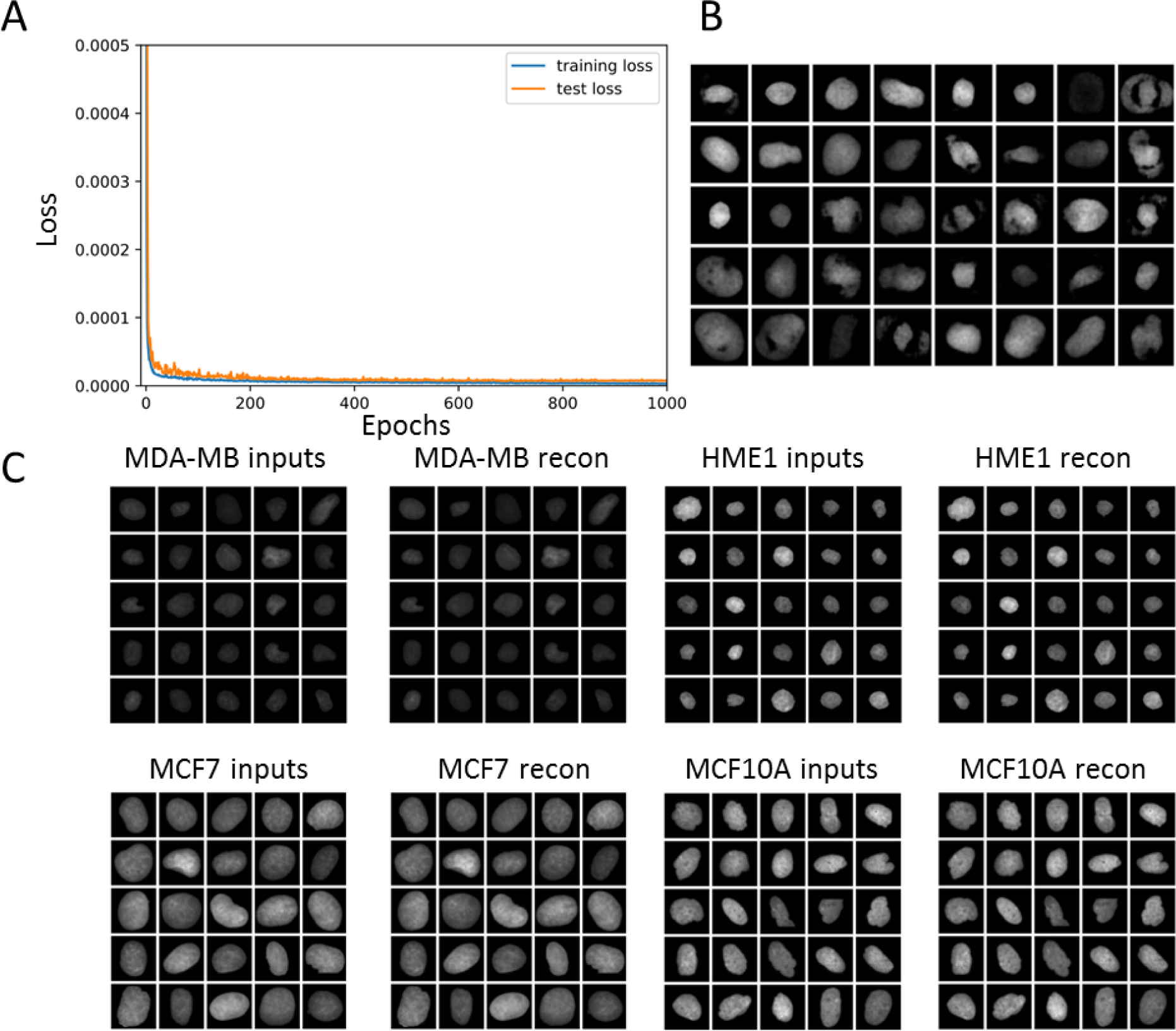
Training the variational autoencoder on various breast cell lines. (a) Training and test loss curves of variational autoencoder plotted over 1000 epochs. (b) Nuclear images generated from sampling random vectors in the latent space and mapped back to the image space. These random samples resemble real nuclei, suggesting that the variational autoencoder learns the image manifold. (c) Input and reconstructed images from different cell lines, illustrating that the latent space captures the main visual feature of the original images.

**Supplementary Figure S16:**
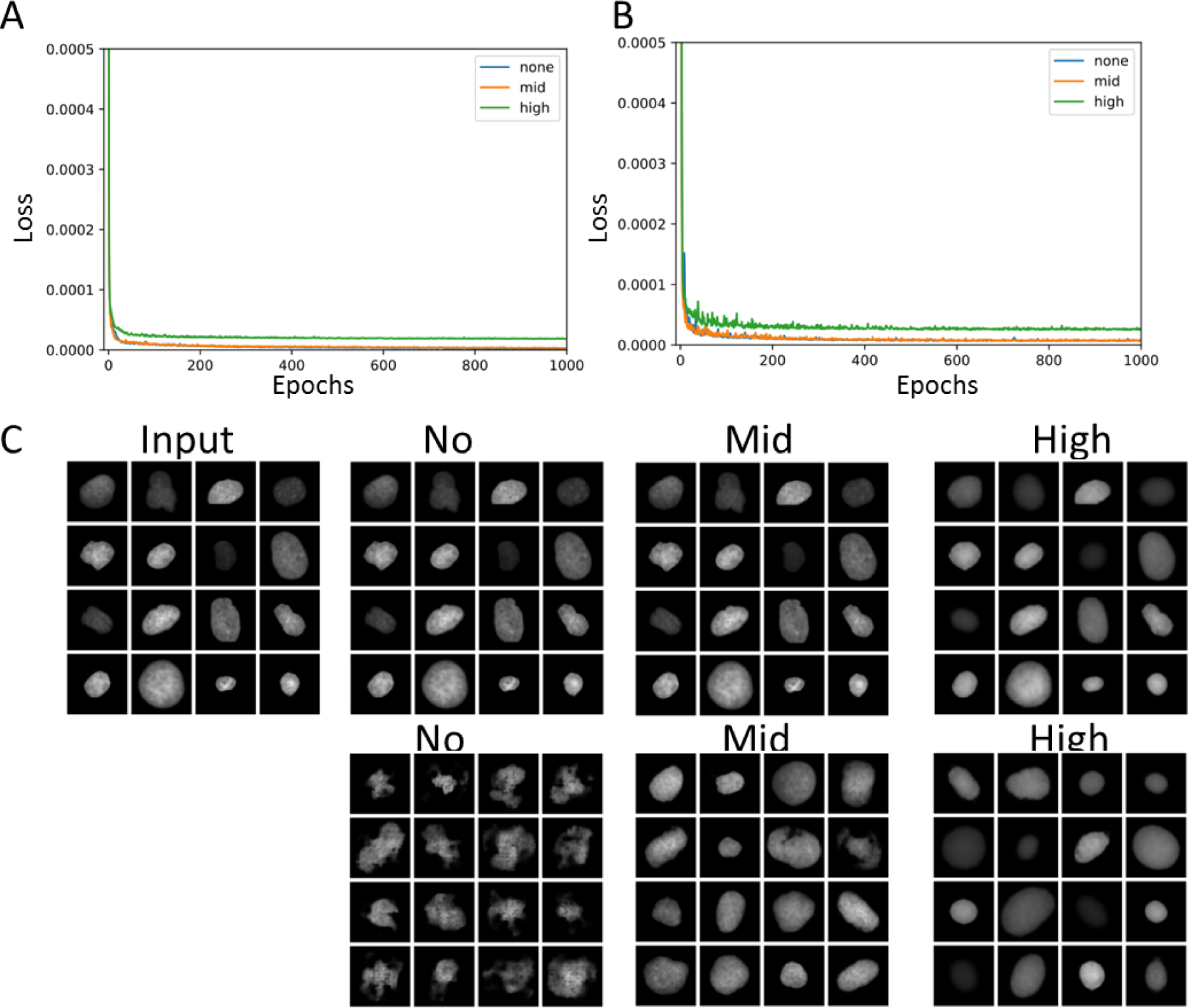
Hyperparameter tuning for variational autoencoder on breast cell lines. (a-b) Training loss and test loss curves respectively with high, mid, and no regularization. (c, top row) Reconstruction results for each model. Models with no or mid-level regularization can reconstruct input images well, while models with high regularization do not. (d, bottom row) Sampling results for each model. Models with no regularization do not generate random samples as well as models with mid-level regularization, which suggests that the model with mid-level regularization best captures the manifold of nuclear images.

**Supplementary Figure S17:**
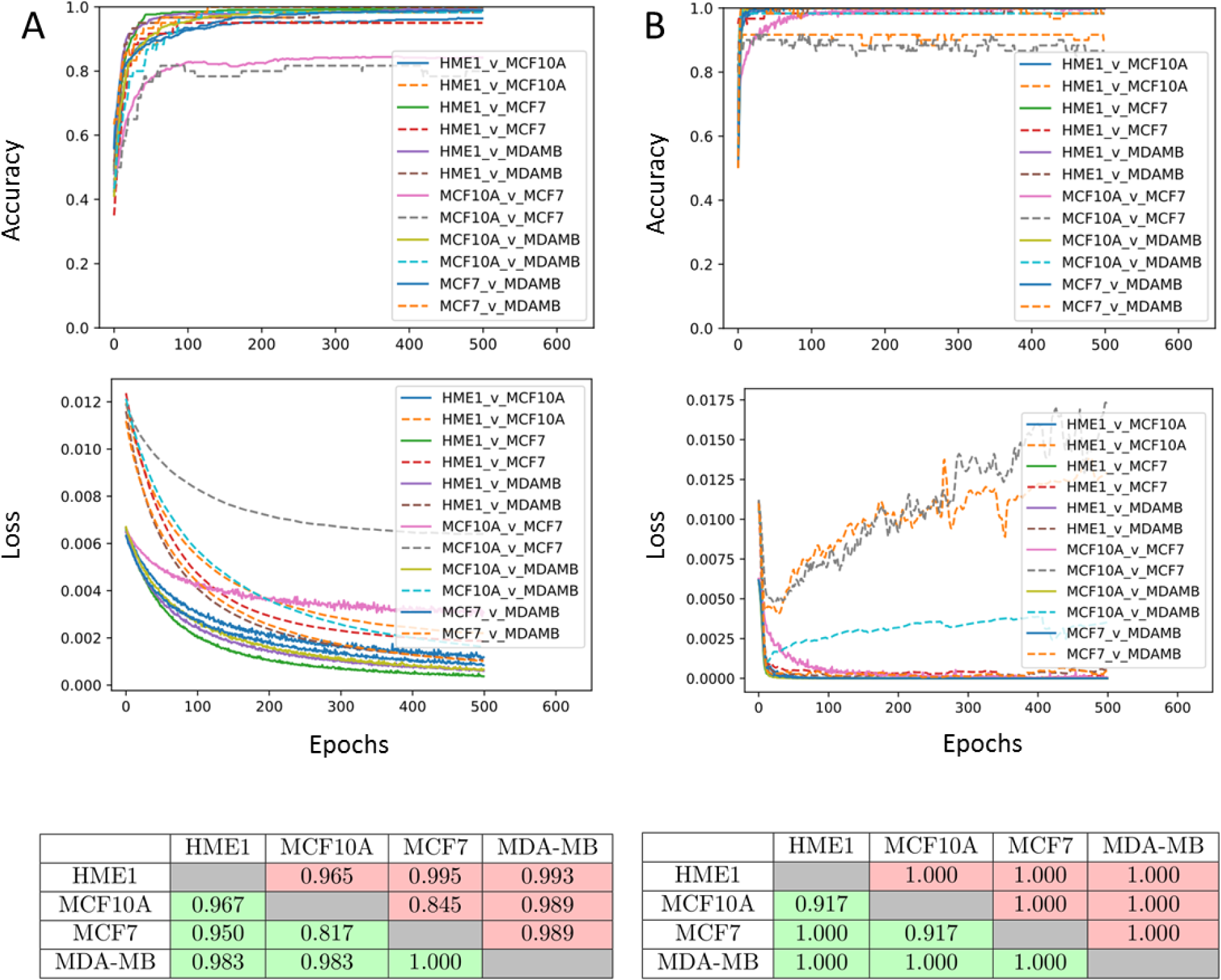
Pairwise classification of nuclei from breast cell lines in the latent space. (a) Classification results in the latent space using a linear model. Top: training and test loss curves for each pairwise comparison (HME1, MCF10A, MCF7, MDA-MB231). Middle: training and test accuracy curves for each pairwise comparison. Bottom: Table of best training (red) and test (green) accuracy for each classification task. (b) Same as (a) but using a two-layer feedforward neural network. For all tasks, the sizes of the training and validation datasets were 550 and 60 respectively.

**Supplementary Figure S18:**
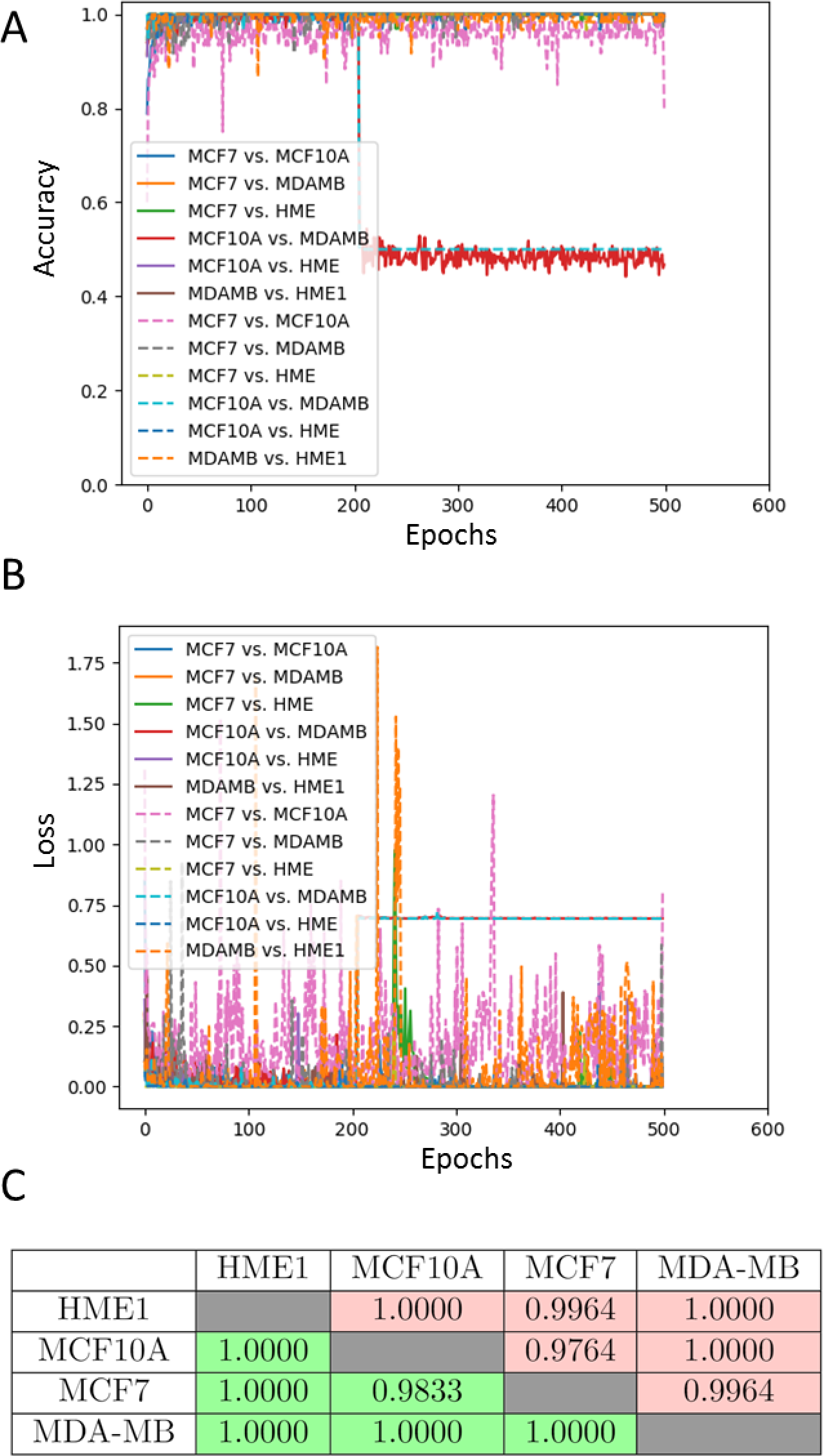
Pairwise classification of nuclei from breast cell lines using VoxNet. (a) Accuracy and (b) loss curves for each pairwise comparison (HME1, MCF10A, MCF7, MDA-MB231). (c) Table of best training (red) and test (green) accuracy for each classification task.

**Supplementary Figure S19:**
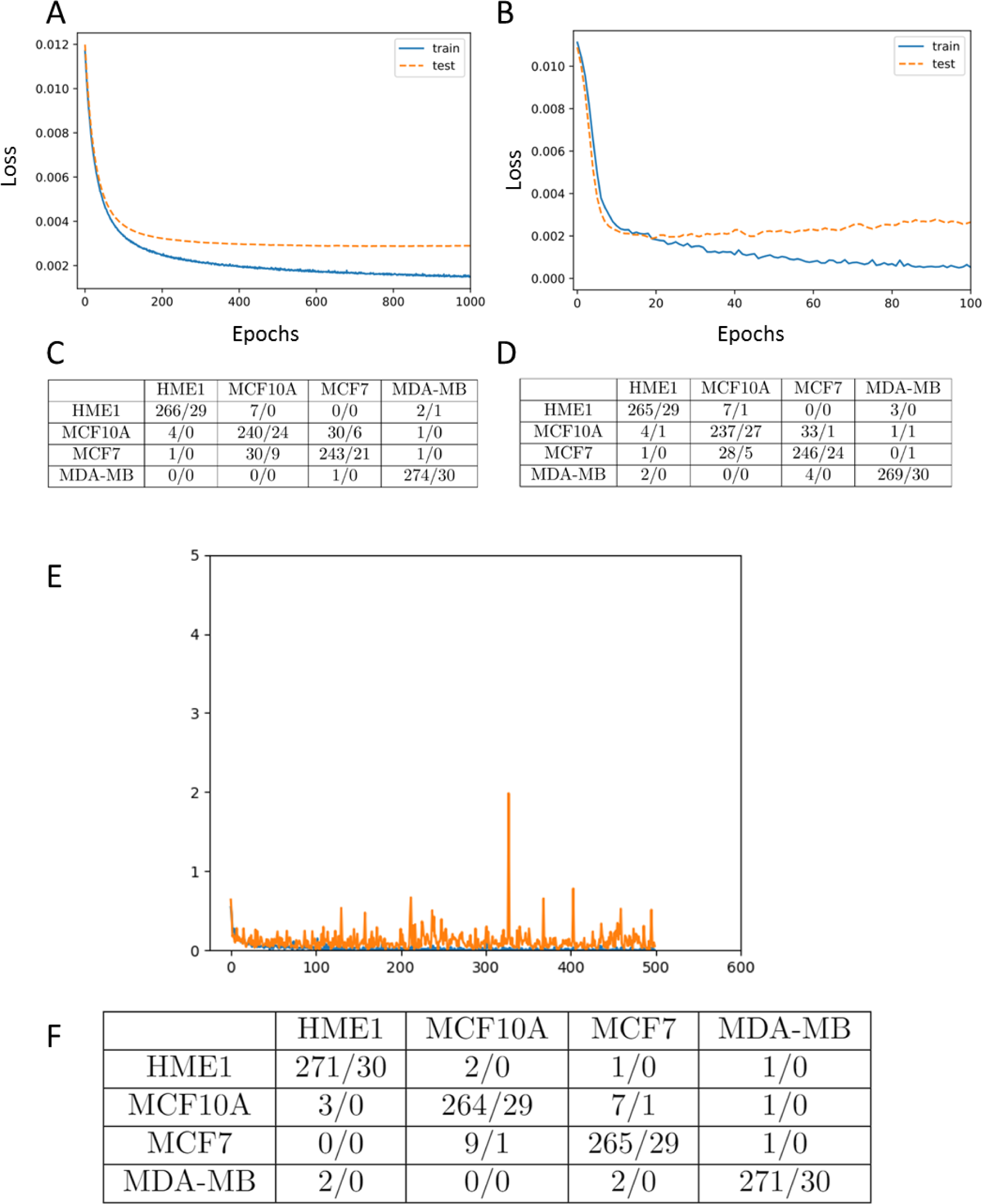
4-way classification results for nuclei from breast cell lines. Training and test loss curves for 4-way classification task of HME-1, MCF10A, MCF7 and MDA-MB231 nuclei using a linear model (a) and 2-layer feedforward neural network (b). (c-d) Confusion matrices for the classification tasks in (a-b). Each entry (X/Y) in row “A” and column “B” indicates that X nuclei of class “A” were classified as “B” in the training set and Y nuclei of class “A” were classified as “B” in the test set. (e-f) Same as (b,d) but for the 4-way classification task in the original image space using a deep convolutional neural network.

**Supplementary Figure S20:**
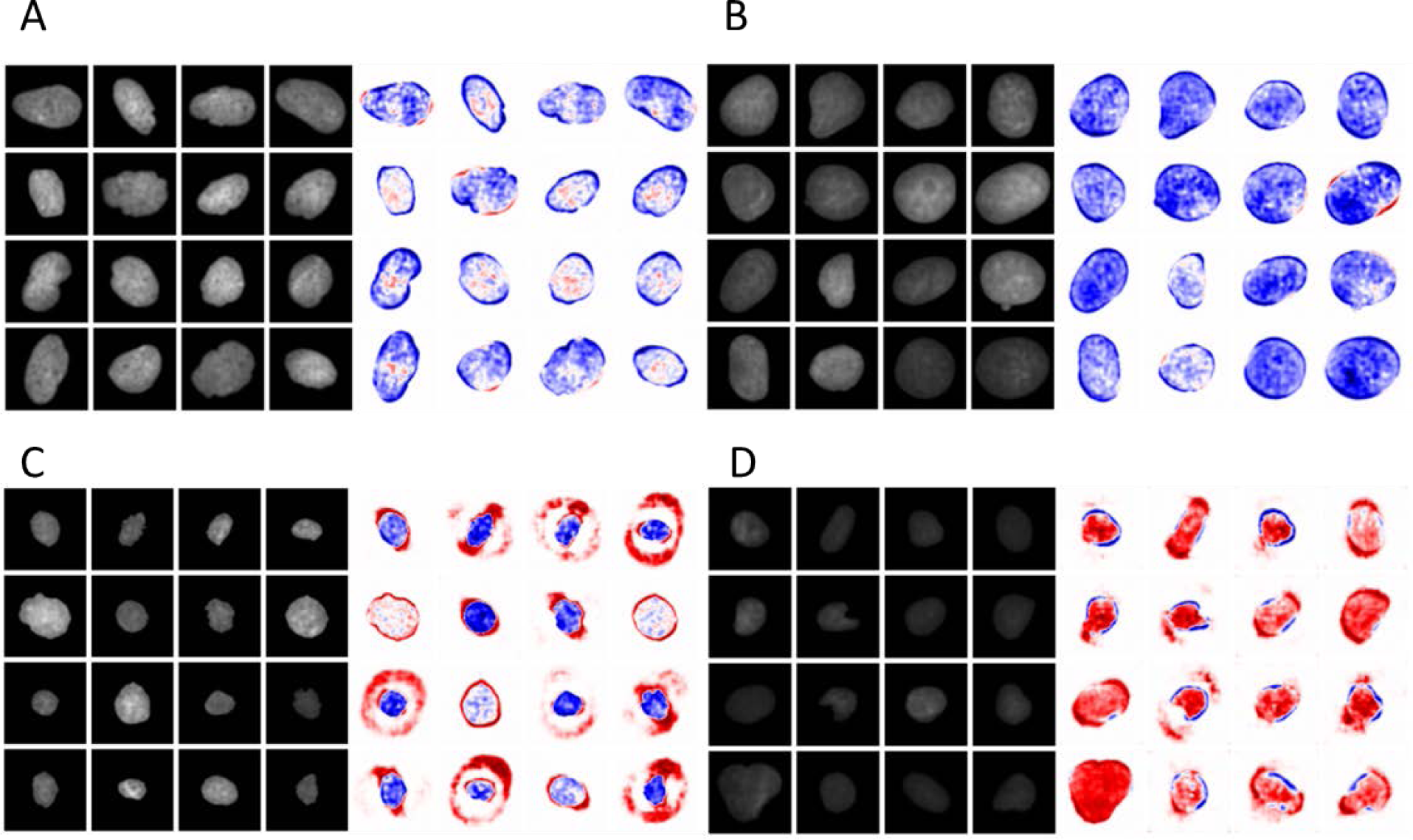
Principal features of change between cell lines. (a) Left: real MCF10A nuclear images. Right: heatmap of changes in pixel intensity of MCF10A nuclei after modulation along the first linear discriminant towards HME-1 nuclei. (b) Left: real MCF7 nuclear images. Right: heatmap of changes in pixel intensity of MFC-7 nuclei after modulation along the first linear discriminant towards MDA-MB231 nuclei. (c) Left: real HME-1 nuclear images. Right: heatmap of changes in pixel intensity of HME-1 nuclei after modulation along the first linear discriminant towards MCF10A nuclei. (d) Left: real MDA-MB231 nuclei images. Right: heatmap of changes in pixel intensity of MDA-MB231 nuclei after modulation along the first linear discriminant towards MCF7 nuclei.

**Supplementary Figure S21:**
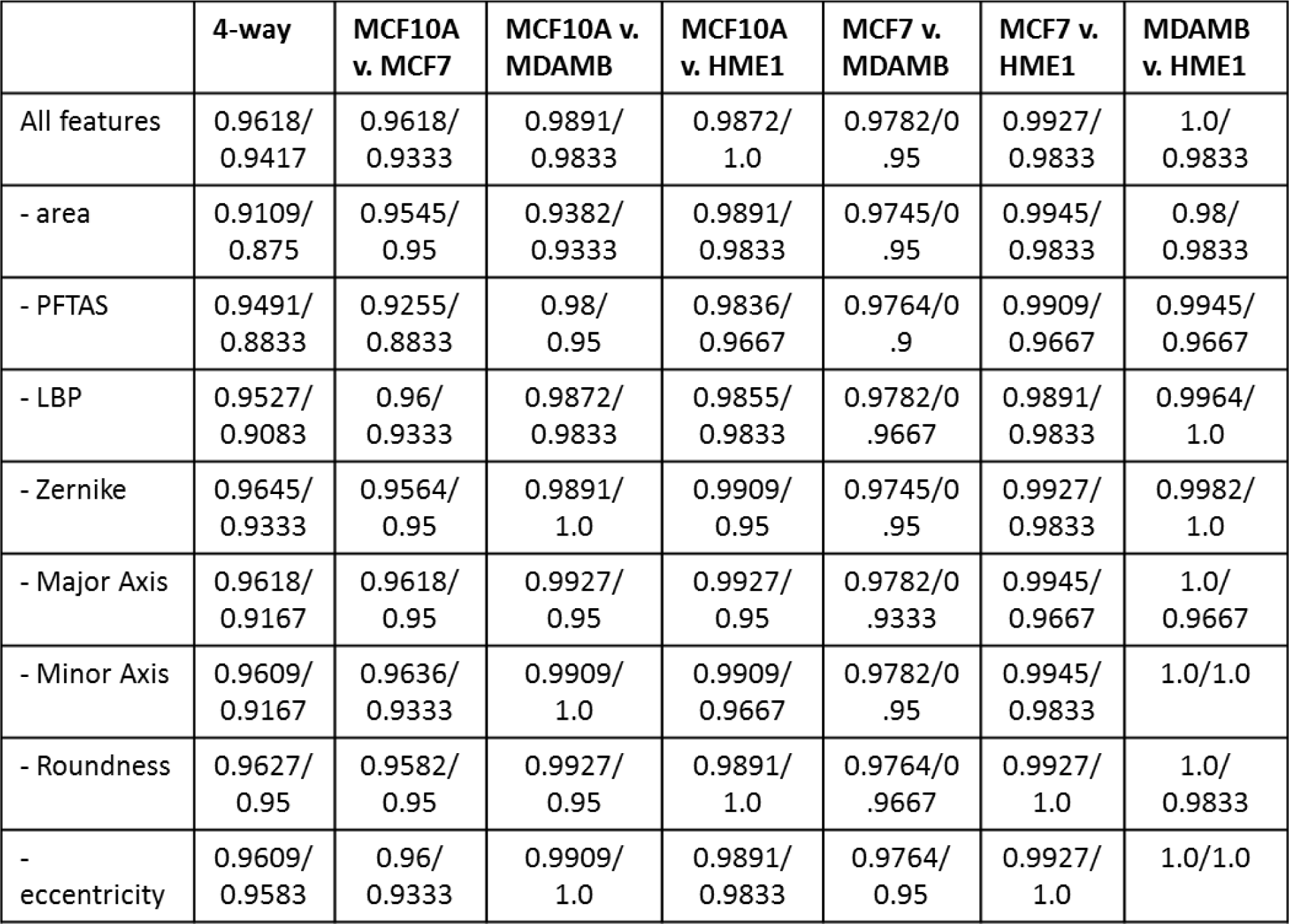
Logistic regression and feature ablation table on the 4 breast cell lines.

**Supplementary Figure S22:**
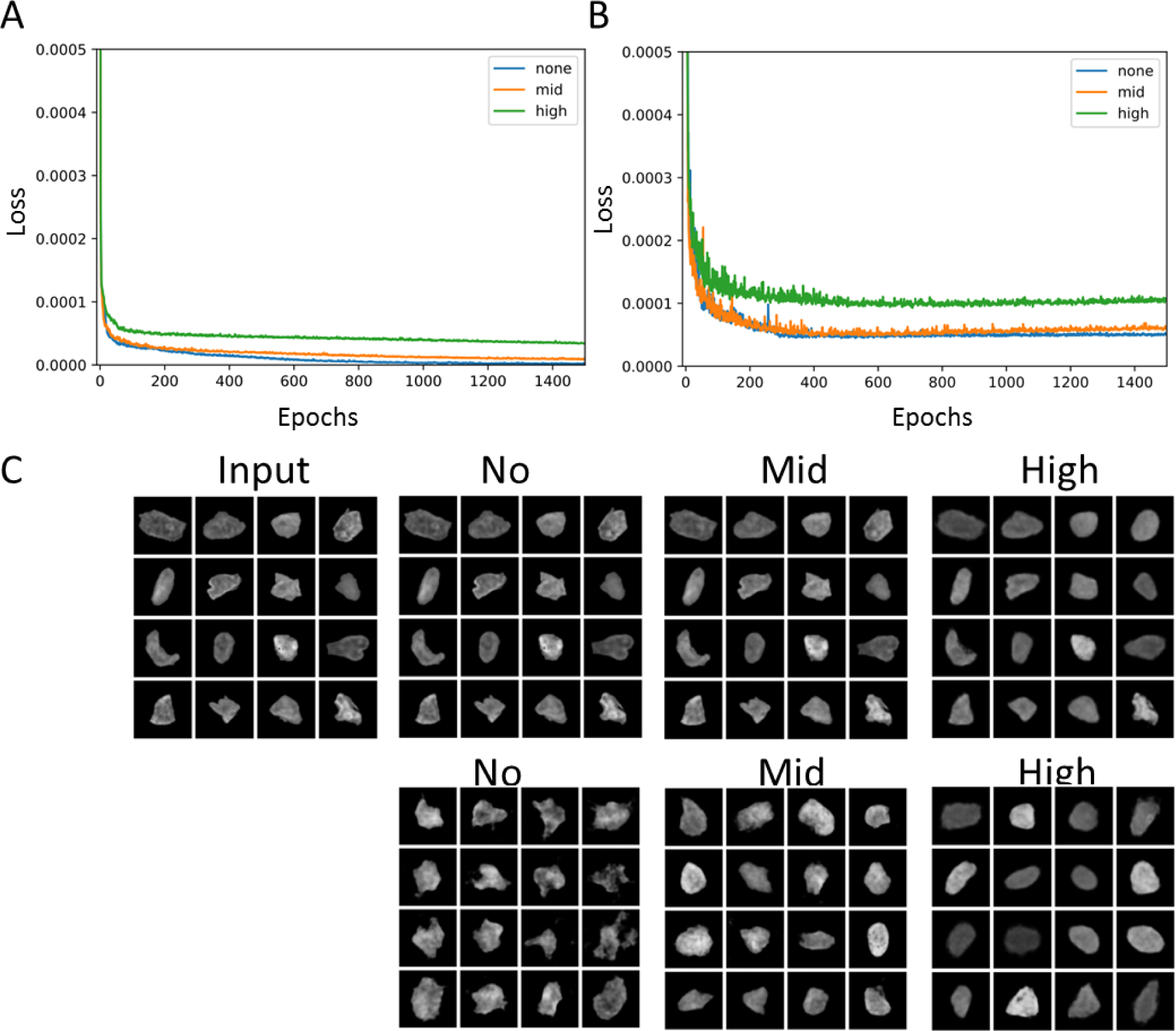
Hyperparameter tuning for variational autoencoder applied to human breast tissues. (a-b) Training loss and test loss curves respectively with high, mid, and no regularization. (c, top row) Reconstruction results for each model. Models with no or midlevel regularization can reconstruct input images well, while models with high regularization do not. (c, bottom row) Sampling results for each model. Models with no regularization do not generate random samples as well as models with mid-level regularization, suggesting that the model with mid-level regularization best captures the image manifold.

